# The imbalance of nature: The Role of Species Environmental Responses for Ecosystem Stability

**DOI:** 10.1101/2025.01.30.635685

**Authors:** Francesco Polazzo, Til Hämmig, Owen L. Petchey, Frank Pennekamp

**Affiliations:** Department of Evolutionary Biology and Environmental Studies, University of Zurich, Zurich, Switzerland

**Author notes:** Corresponding author: Francesco Polazzo. The two authors contributed equally to this work (Francesco Polazzo, Til Hämmig). Joint senior authorship (Owen L. Petchey, Frank Pennekamp). Author contributions: FP: Conceptualisation, Formal analysis, Investigation, Methodology Writing - original draft, Project administration, Visualisation, Software. TH: Investigation, Data curation, Methodology, Writing - review & editing, Visualisation, Software. OLW: Conceptualisation, Funding acquisition, Methodology, Project administration, Resources, Supervision, Writing - review & editing. FP: Conceptualisation, Methodology, Project administration, Supervision, Writing - review & editing. Open Science Statement: Data and code to reproduce the analysis is available at https://github.com/FrancescoPola/Response-div-stability-competitive-communities/tree/main, and will be permanently deposited in Zenodo upon publication.

## Abstract

Understanding the mechanisms underlying ecosystem stability is crucial in predicting ecological responses to environmental fluctuations. While the diversity-stability relationship has been widely studied, the role of species’ fundamental responses to the environment remains underexplored. Here, we investigate how the distribution of fundamental responses, captured by a novel metric—imbalance—drives ecosystem stability through asynchrony and population stability. Using a microcosm experiment with protist communities, we manipulated species richness and response distributions (defined as interspecific variation in species performance curves) under fluctuating temperature and different nutrient concentrations. Our results show that lower imbalance, achieved through asynchrony or high population stability, causes higher temporal stability, while richness has no effect on stability. Structural equation modelling revealed that imbalance decreases stability indirectly via increasing synchrony and decreasing population stability, explaining 90% of observed variation. Comparing imbalance derived from single versus multispecies communities demonstrates that fundamental species responses are primary drivers of stability, challenging traditional paradigms emphasizing interspecific interactions. This study provides mechanistic links between species’ responses, environmental variability, and ecosystem stability, offering new insights into the responses of ecological systems to environmental change.

## Introduction

The diversity-stability debate has been a cornerstone of ecological research for decades ^1^. The question, “Does diversity promote stability, and how?” remains unresolved, as both diversity ^2^ and stability ^3^ are complex, multidimensional concepts. While many studies report a positive relationship between species diversity and the temporal stability of ecosystem properties ^4–6^, some studies document negative relationships ^7,8^. Reconciling these contrasting results is challenging, given the complexity of mechanisms underlying the diversity-stability relationship.

One proposed mechanism is that diversity leads to higher temporal stability of community and ecosystem properties due to the greater range of species’ fundamental responses to environmental change expected in more diverse communities ^9,10^. A more diverse community is likely to exhibit greater response diversity, whereby species vary in their reactions to environmental fluctuations ^11^. High response diversity occurs when some species increase in abundance while others decrease in response to environmental changes ^12^. In fluctuating environments, response diversity should promote asynchrony in species’ abundance over time, a key stabilizing mechanism according to the insurance hypothesis ^13^.

The conceptual foundation linking response diversity to stability is well-established ^9,11,14^.

Yet, high ecosystem stability can also arise without high response diversity when all species exhibit relatively weak responses to environmental change ^15^. In such cases, population stability—rather than asynchrony—underpins ecosystem stability, as weak responses ensure low variability across species. Thus, fundamental species’ responses can influence ecosystem stability through two complementary mechanisms: (1) by affecting asynchrony via response diversity and (2) by affecting population stability. These mechanisms interact to shape ecosystem stability, with their combined effects determined by the ensemble of species’ fundamental responses that occur in a community ^16^.

Despite the solid theoretical foundation of this framework, controlled experiments explicitly testing the role of fundamental species’ responses in driving stability remain scarce ^17,18^. To address this gap, we performed a large microcosm experiment involving aquatic protist communities. We factorially manipulated species richness and the distribution of species’ fundamental responses under fluctuating temperature regimes across three levels of nutrient availability. To obtain robust estimates of species’ intrinsic growth rate (our measure of species’ fundamental responses), we estimated response surfaces of six species across all combinations of five temperature levels and five nutrient concentrations ^19^ in an independent experiment. Communities of 2, 3, or 4 species were then assembled. Within each richness level there were different compositions, in order to manipulate the distribution of species’ fundamental responses. These ranged from distributions closely centred around zero, reflecting a mix of positive and negative responses with similar magnitudes, to skewed, with uneven response magnitudes or predominantly unidirectional responses. Communities were then subjected to fluctuating temperature regimes and three constant nutrient conditions, and ecosystem stability was measured to answer our primary question: How do species’ fundamental responses determine ecosystem stability?

We predicted that communities with distributions of species’ responses to environmental change centred around zero (i.e., balanced communities), characterized by a mix of positive and negative responses with similar magnitudes (i.e. high response diversity) or small overall responses (i.e. high population stability), would exhibit greater stability. In contrast, communities with predominantly positive or negative species’ responses (i.e., imbalanced communities) were expected to be less stable.

To quantify these distributions, we introduced a new metric, *imbalance* (Fig. 1), which captures the symmetry and magnitude of species’ responses around zero. Imbalance is low when the distribution of species’ responses is centred on zero, thus reflecting both asynchrony and population stability ^16^. We expect low values of imbalance to be associated with high ecosystem stability. This is, a balanced system should be a stable system.

**Figure 1.**
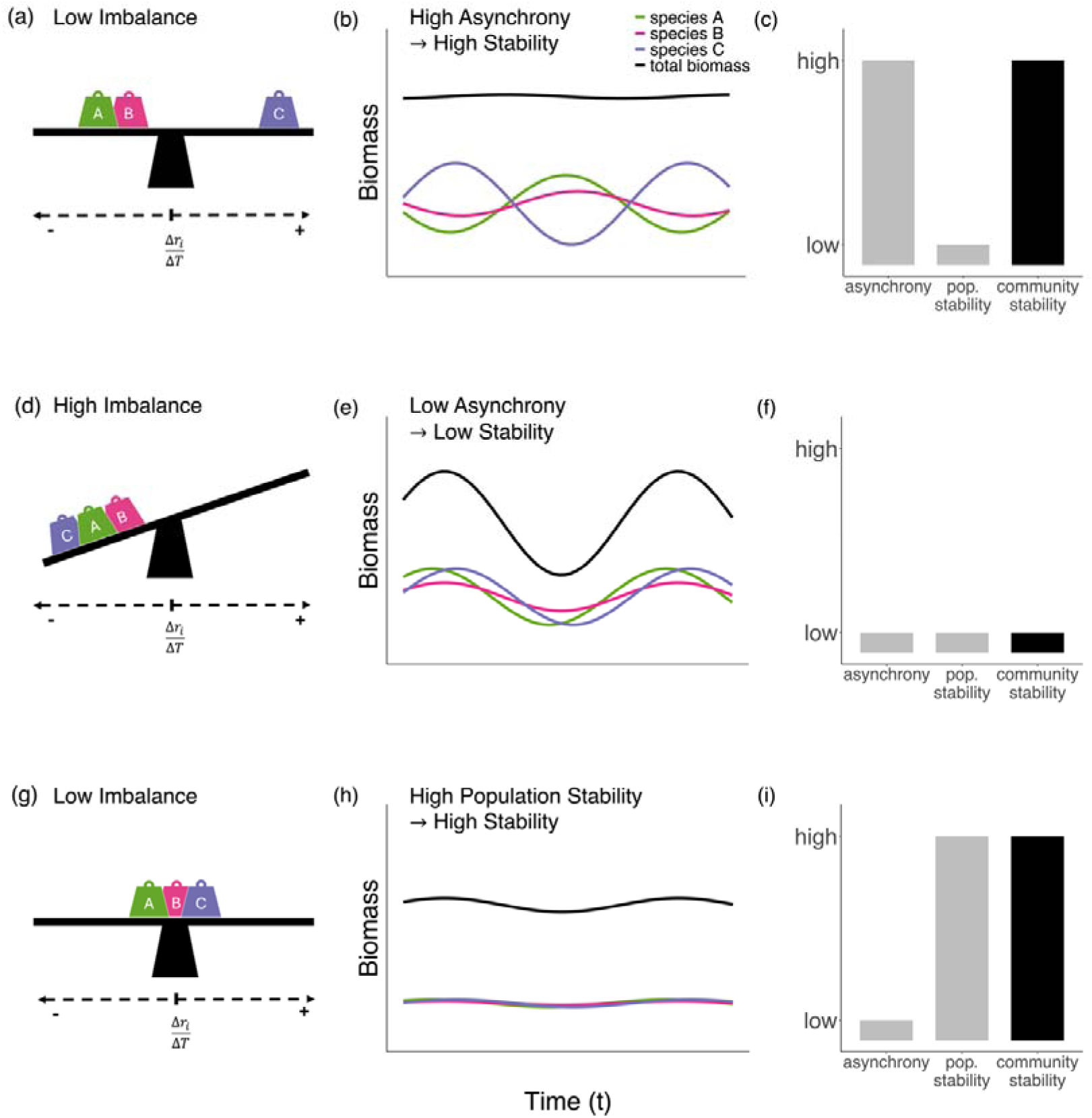
Conceptual illustration of imbalance driving community stability through response diversity and population stability. Panels (a–c) depict a balanced system with high response diversity. (a) Species A and B respond negatively to the temperature change, while species C responds positively. (b) Asynchronous biomass fluctuations ensure total community biomass varies little over time. (c) High community stability results from high asynchrony and low population stability. Panels (d–f) depict an imbalanced system with low response diversity. (d) Species A, B, and C respond similarly to the temperature change. (e) Synchronous biomass fluctuations result in greater variation in total community biomass over time. (f) Low community stability arises from low asynchrony and low population stability. Panels (g–i) illustrate a balanced system with high population stability. (g) Species A, B, and C respond similarly, but very weakly, which results in stable population dynamics. (h) High population stability leads to stable community biomass over time. (i) High population stability, despite low asynchrony, results in reduced community stability.

Our prediction that species’ responses from monocultures predict community stability carries the implicit assumption that other mechanisms, such as interspecific interactions ^20,21^ are expected to be relatively unimportant. Furthermore, by manipulating response distributions within each richness level, we minimized effects of species richness such as statistical averaging (i.e. portfolio effect). In order to test if the effects of species interactions could be important, we calculated two types of imbalance: fundamental imbalance and realised imbalance. In analogy to the fundamental niche, *fundamental* imbalance is based on species fundamental responses to an environmental change. In *realized* imbalance, fundamental responses were weighted by the realized relative species biomass of species during the community experiment. Comparing the explanatory power of fundamental and realized imbalance allowed us to compare how the interactions between species in a community influence stability. This comparison provides insight into the role of species interactions in driving community stability, highlighting whether interspecific interactions contribute to stability by buffering or enhancing the effects of environmental changes, or whether the observed stability is primarily a result of the fundamental responses of species without significant interaction effects.

This study tests for evidence of two biological mechanisms underpinning the diversity-stability relationship. By explicitly manipulating the distribution of species’ fundamental responses we move beyond correlative approaches that infer mechanisms from patterns (e.g. high diversity = greater asynchrony). Our approach is novel in three ways: (1) it isolates the effect of fundamental response distributions, controlling for confounding factors and demonstrating the robustness of these mechanisms across different richness levels and environmental contexts; (2) it assesses the roles of asynchrony and population stability in driving ecosystem stability; and (3) it provides a new approach to understand the link between species’ fundamental responses and ecosystem stability.

## Results

In our experiment, we found that greater imbalance was associated with lower stability (Estimate = -0.08, SE = 0.02, p < 0.001; Fig. 2). The difference between realized (weighted by species biomass contribution) and fundamental (only based on species’ responses measured in monoculture) imbalance (Fig. 2) was not significant (χ² = 0.49, df = 1, p = 0.48). We thus use fundamental balance for all further analyses.

**Figure 2.**
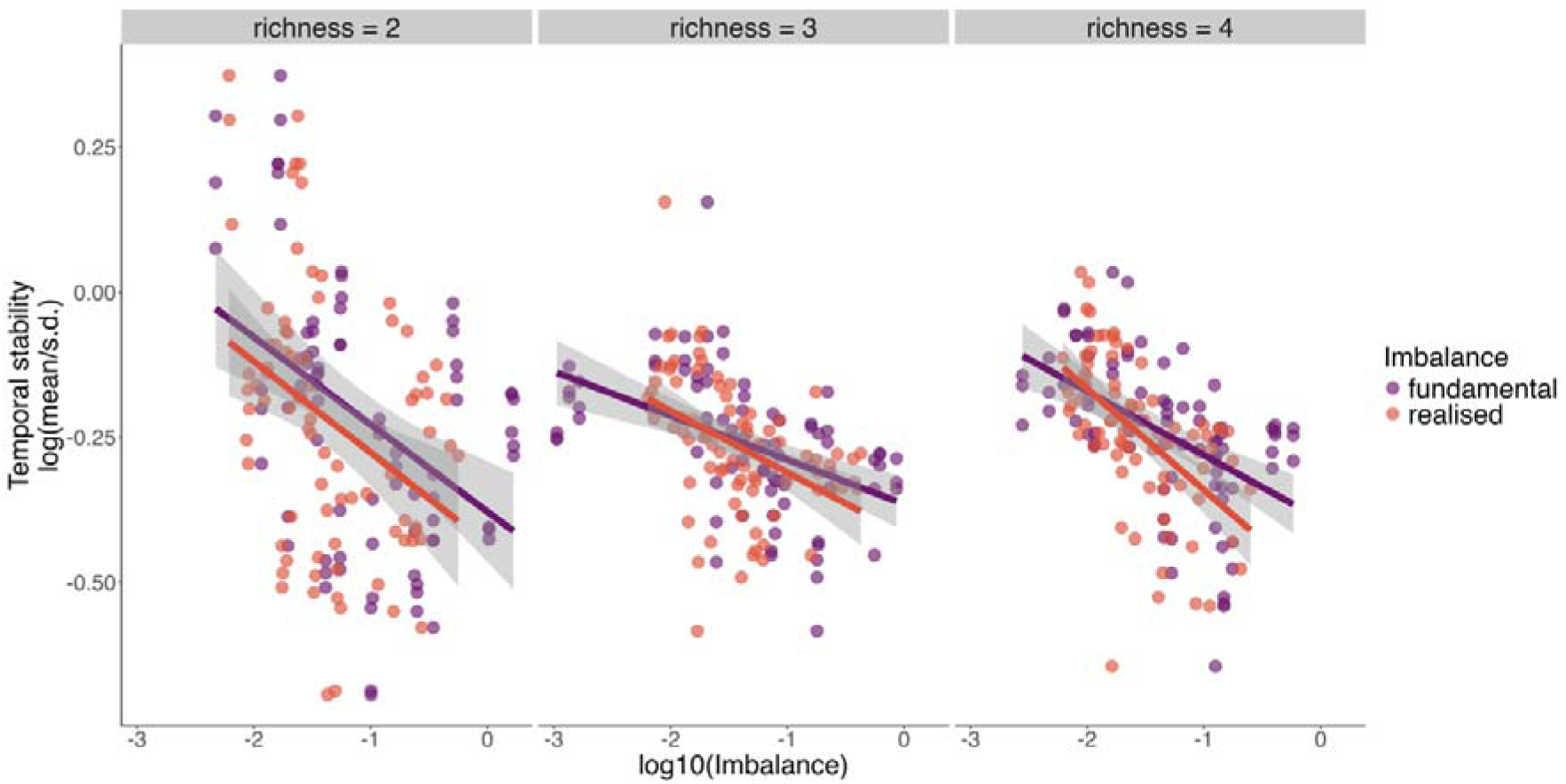
Effects of fundamental and realised imbalance on total community biomass temporal stability (n = 243).

Richness did not affect stability (Fig. 3a). Nutrient concentration had a positive effect on stability (estimate = 0.25, SE = 0.01, p < 0.001; Fig. 3c), whereas temperature had a destabilizing effect (estimate = -0.10, SE = 0.02, p < 0.001; Fig. 3b). Finally, temperature and nutrients interacted non-additively, reducing stability (estimate = -0.09, SE = 0.02, p = 0.004).

**Figure 3.**
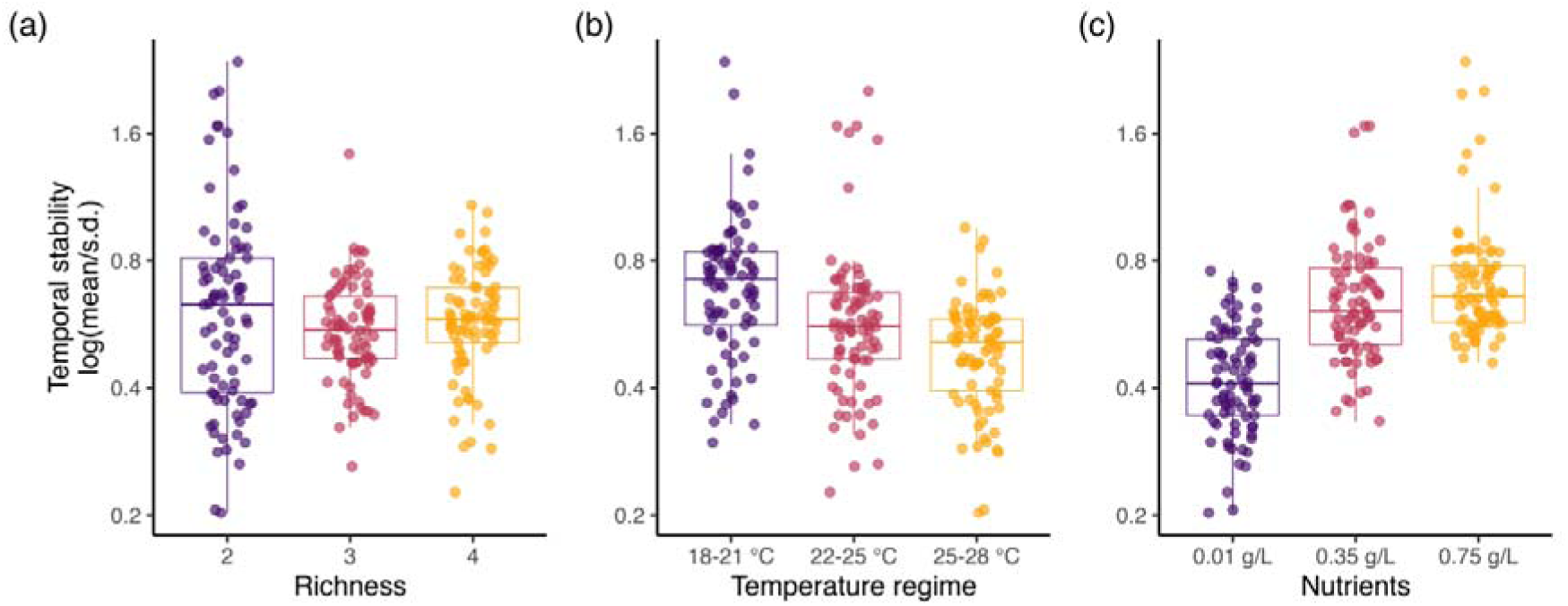
Effects of richness (a), temperature (b), and nutrients (c) on community total biomass temporal stability (n = 243).

We then used a structural equation model (SEM) to understand how imbalance influences stability via asynchrony (Fig. 4a and b) and population stability (Fig. 4c and d). An excellent model fit to the observed variance-covariance structure was obtained (Chi square = 2.36, p = 0.50; CFI = 1.0, RMSEA = 0.00, Fig. 5). The SEM shows that imbalance had a negative direct effect on asynchrony (standardized coefficient = - 0.18, p < 0.001) and on population stability (standardized coefficient = - 0.30, p = 0.006). Both asynchrony (standardized coefficient = 0.33, p < 0.001) and population stability (standardized coefficient = 0.99, p < 0.001) had a positive effect on temporal stability, jointly explaining 90% of the variation in temporal stability in our experiment (Supplementary Information 1, Section 10). Nutrient concentration had a negative effect on asynchrony (standardized coefficient = -0.50) and a positive effect on population stability (standardized coefficient = 0.65).

**Figure 4:**
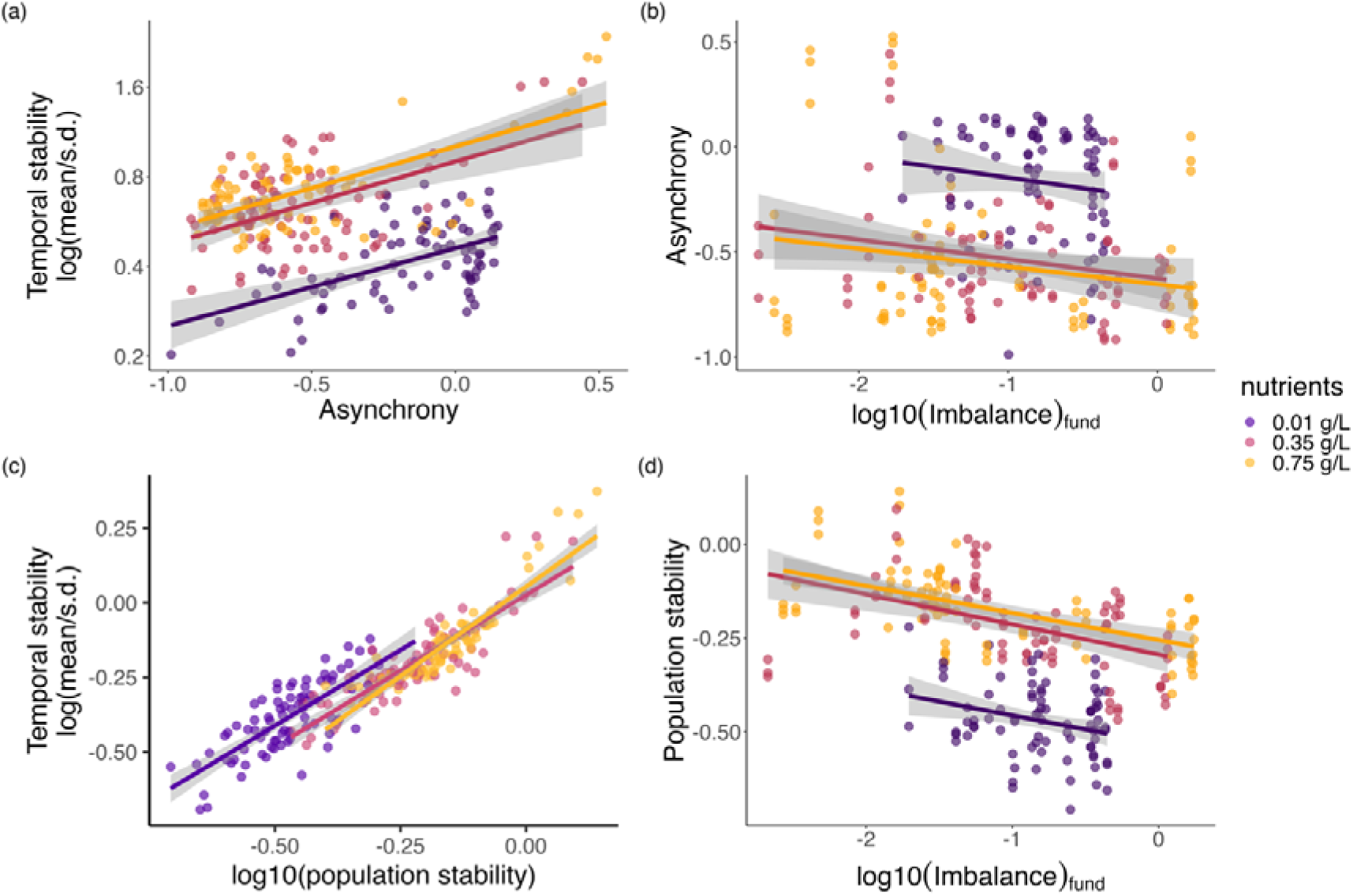
(a) Relationship between temporal stability and asynchrony (Gross) divided by nutrient level. (b) Relationship between asynchrony (Gross) and imbalance divided by nutrient level. (c) Relationship between log10 of population stability and log 10 of ecosystem stability.

**Figure 5.**
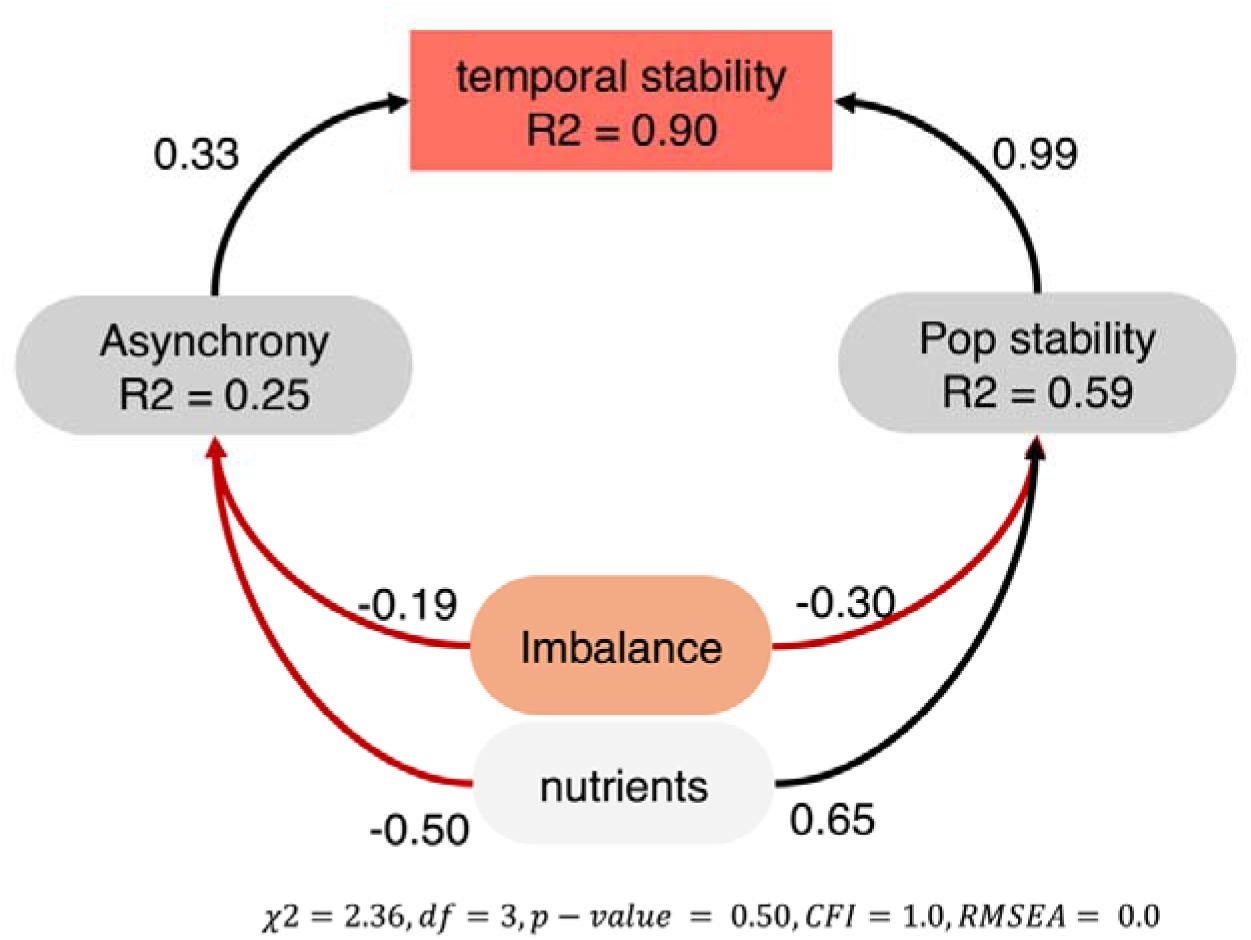
Structural Equation Model (SEM) showing the direct effect of imbalance and nutrients on asynchrony and population stability, and direct effects of asynchrony and population stability on community temporal stability. Numbers on the arrows indicate the standardized coefficients. Negative effects are indicated as red arrows, whereas positive effects are represented as black arrows. All effects are significant. The full table of coefficients is displayed in the Supplementary Information 1, Section 10.

## Discussion

Our study demonstrates that the fundamental responses of species to environmental fluctuations play a critical role in driving ecosystem stability, mediated through two complementary mechanisms: asynchrony and population stability. By introducing the novel metric of imbalance, we showed that balanced distributions of species’ responses—whether through high response diversity or weak responses—enhance temporal stability in community biomass. This challenges traditional views emphasizing interspecific competition as the dominant stabilizing mechanism ^10,21,22^ and aligns with recent findings in natural ecosystems where environmental variability outweighs competitive effects ^18,23^. Importantly, we found that while species richness was not associated with stability, the distribution and magnitude of species’ fundamental responses to nutrient availability and temperature, were key determinants. These findings provide a mechanistic understanding of the diversity-stability relationship, advancing our ability to predict ecosystem dynamics in fluctuating environments.

When response diversity is high, differences in how species respond to environmental fluctuations can promote asynchrony in population dynamics, reducing temporal variability at the community and ecosystem level ^9,14,18^. In contrast, when species exhibit minimal responses to environmental changes, population stability directly contributes to ecosystem stability, even in the absence of asynchrony ^16,24^. Our findings highlight the dual importance of asynchrony and population stability but also suggest that species’ fundamental responses are the dominant mechanism driving asynchrony. The extent to which species’ fundamental responses drive positive diversity-stability relationships has been debated ^10,20,21^. Several alternative mechanisms have been proposed, including compensatory dynamics arising from interspecific competition ^20^, evenness of mean abundance ^25^, overyielding ^26^, and statistical averaging effects ^27^. However, our results systematically argue against these alternative mechanisms operating strongly in our study: Minimal differences between fundamental and realized imbalance suggest that species interactions played a limited role, as compensatory dynamics arising from competition would have caused larger discrepancies. The limited role of species interactions in determining stability was further supported by a direct quantification of species interaction coefficients (Supplementary Information 3). High species evenness in biomass contributions in our study (Supplementary Information 1, Fig. 11) further reduces the potential for selection effects, where stability is driven by dominant species ^23^. However, this finding does not rule out that evenness may be important in other experiments, particularly when uneven biomass distribution can increase the importance of dominant species ^28^. Stability in our communities emerged largely from species’ fundamental responses to environmental change. Additionally, we did not find any effect of species richness, suggesting that statistical averaging (i.e. portfolio effect) did not play any role in stabilizing community biomass. These results support the hypothesis that stability is primarily driven by response diversity promoting asynchrony and from population stability arising from weak species responses, consistent with theoretical predictions ^9,16^. We also hypothesize that the larger role of population stability in driving community stability relates to the weak response species showed to the experimental temperature fluctuations. This may be related to the temperature fluctuations being within the thermal niche of the species involved in this experiment, which prevented large fluctuations at the population level, and thus potentially limiting the role of asynchrony. Whether the distribution of species’ responses (e.g., as quantified by imbalance) is an important mechanism driving the diversity stability relationship in other ecosystems and environmental setting, for example where species interactions may play a stronger role remains a major ecological question.

Furthermore, we found a significant and direct effect of temperature, nutrients, and their interaction on stability. High temperatures tended to reduce response diversity by synchronizing species’ responses, increasing imbalance, and destabilizing communities. On the other hand, higher nutrient levels stabilized total biomass. However, at lower nutrient levels, higher temperature fluctuations can promote response diversity, fostering stability. The interaction between temperature and nutrients amplifies these effects, with high temperatures and nutrients leading to less balanced species responses and lower stability.

We hypothesised that the additional effect of the environmental conditions on stability not captured by imbalance could be explained by the fact that we used fundamental imbalance as predictor of stability. While fundamental imbalance was calculated from monocultures exposed to constant temperature and nutrients, our community experiment featured fluctuating temperatures. The realization that the effect of environmental fluctuations can have biological consequences that cannot be inferred from the effect of mean environmental conditions is formalized by Jensen’s inequality ^29^. In summary, while fundamental species responses are key to understanding stability, the environment remains a dominant factor in shaping community stability in fluctuating conditions.

We quantified the distribution of species’ responses using a new metric, imbalance, that allowed us to predict ecosystem stability from fundamental information about species (i.e., information collected while the species were in isolation from other species). Previous methods for measuring the distribution of species’ responses have focused on response diversity, and recently proposed metrics to summarise the diversity of species responses in a community include dissimilarity and divergence ^12,30^. While these metrics are useful, they have limitations. For example, dissimilarity captures the total variation in species’ responses but does not account for the direction (sign) of the responses, while divergence only considers the responses of the two most extreme species. Furthermore, high values of divergence are possible even when all species have very weak responses to environmental change (i.e., all species are very stable). Conversely, imbalance quantifies the symmetry around zero of all species’ responses accounting for their magnitude. Critically, imbalance systematically outperformed divergence in predicting the relationship between the distribution of species’ responses and ecosystem stability (Supplementary Information 1,

Section 5). Moreover, while divergence and dissimilarity are limited to assessing response diversity ^12^, imbalance captures both response diversity and population stability. By capturing these two fundamental and interlinked mechanisms driving ecosystem stability ^16,31^, imbalance provides a more robust and integrative measure of the factors underpinning ecosystem stability.

Our findings explain why increased species richness will not always enhance community stability and therefore have the potential to explain the variation in effects of species richness on ecosystem stability reported in the literature. A negative relationship between diversity and stability ^7,8^ may arise when increased richness does not result in higher response diversity nor population stability, thereby providing no stabilizing effect. This highlights the importance of considering both the distribution of species’ responses and their dependence on environmental variability when evaluating the diversity-stability relationship.

By integrating the distribution of species’ responses into stability predictions, we offer a robust approach for understanding community dynamics in fluctuating environments. This framework emphasizes that stability is not merely a consequence of species richness but depends on the balance of species’ responses and their relationship to environmental variability. Our experimental results underscore the critical role of environmental variability in shaping ecosystem stability, providing new insights into the mechanisms that link diversity, environmental drivers, and ecosystem stability.

## Methods

### Response Surface Experiments

To assemble communities with varying distributions of species’ responses to environmental change, we first determined intrinsic growth rate response surfaces for six ciliate species (*Colpidium striatum*, *Dexiostoma campylum*, *Loxocephalus* sp., *Paramecium caudatum*, *Spirostomum teres*, and *Tetrahymena thermophila*). Each species was grown in monoculture under a fully factorial design comprising five nutrient levels and five temperature treatments (25 treatments total).

Long-term stock cultures were maintained at 15 °C in bacterised alfalfa medium (0.55 g/L) with 0.5 mL/L of Chalkley’s medium (10X). For the experiment, we prepared five alfalfa concentrations (0.01 g/L, 0.75 g/L, 1.25 g/L, 2.5 g/L, 3.0 g/L) and incubated the cultures at constant temperatures (18, 21, 24, 26, or 28 °C) in programmable incubators (IPP260plus, Memmert, Germany). Each treatment had three replicates, resulting in 450 microcosms (50-mL Sarstedt tubes). Microcosms were initiated with 25 mL of medium, with a starting density of 100 individuals/mL for most species, except *Spirostomum* and *Paramecium*, which were started at 50 individuals/mL. Over three weeks, population densities were measured daily for the first five days and every other day thereafter (4’950 unique samplings). Medium removed during sampling was replaced with fresh sterile medium with the same nutrient concentration.

Before terminating the experiment, we visually inspected the measured cell density time-series to assure that populations had reached the stationary growth phase.

Initially, we planned to assemble competitive communities from a pool of six ciliate species. However, *Tetrahymena thermophila* grew poorly in several treatments preventing us to reliably estimate growth rates (Supplementary Information 1, Fig. 1). Therefore, we decided to exclude it from the community experiment. For the remaining five species, densities and dynamics were consistent with prior experiments ^32,33^, and were thus retained for further analysis.

Intrinsic growth rates were estimated from monoculture data by regressing log-transformed population densities against time during the exponential growth phase ^34^.

The exponential phase varied among species and treatments (Supplementary Information 2, Fig. 1); for instance, we used data up to day 8 for *Dexiostoma* and *Paramecium*, and the first three days for *Colpidium* and *Loxocephalus* at 28 °C. *Spirostomum* grew much slower than all the other species, so its entire time series was used. Growth rate surfaces were modelled using generalized additive models (GAMs) with temperature and nutrients as predictors, employing the *mgcv* package in R ^35^, because they allow for non-linearity and interactions between environmental variables (i.e. temperature and nutrients) by making use of tensor-product smooths ^30^.

### Community experiment

To test the role of species’ fundamental responses for the temporal stability of community biomass, we conducted a community experiment under varying environmental contexts. Our environmental treatments involved three nutrient levels and three temperature fluctuation regimes. While nutrient concentrations remained constant through time, temperature alternated between two values in each regime, with three days at one temperature followed by a one-day transition to the next. The three different temperature fluctuation regimes were: 18–21 °C, 22–25 °C, and 25–28 °C, with fluctuations maintained using programmable incubators.

We designed temperature regimes informed by the species-specific response surfaces generated during the response surface experiments. Using this information, we selected temperature regimes that systematically varied in their degree of response divergence (see next section), with the goal of capturing a gradient of fundamental species responses within each regime. We maintained each temperature constant for three days before transitioning to the next to ensure sufficient time for differences in growth rates at varying temperatures to manifest.

We used bacterised alfalfa medium (0.01 g/L, 0.35 g/L, or 0.75 g/L) as in the response surface experiment. Resource species (*Bacillus subtilis*, *Serratia fonticola*, and *Brevibacillus brevis*) were cultured separately in DSMZ-1 medium at 37 °C for 24 hours, then combined after three days of growth in alfalfa medium. Communities were initiated using stocks grown at 22 °C, slightly below the mean temperature range. We introduced each species at 10% of its mean carrying capacity (based on data from the response surface experiment), keeping the total microcosm volume fixed at 25 mL.

### Community assembly

To explore the relationship between species’ fundamental responses and stability, we aimed to assemble communities that spanned a gradient of species’ responses distribution while varying species richness. Specifically, we selected community compositions that allowed us to test how differences in responses distribution influence stability across multiple environmental conditions.

We assembled communities with 2, 3, or 4 species from a pool of 5 species, creating all possible community compositions at each richness level. To ensure a representative gradient of responses distribution within each richness level and environmental treatment, we quantified the distributions of species’ responses for all possible community compositions.

We then selected three compositions per richness level that captured a range of response divergence values, including distributions where species’ responses were centred around zero, strongly skewed (dominated by positive or negative responses), and intermediate cases where responses were neither perfectly centred nor highly skewed. To quantify response divergence, we used the following metric ^12^:

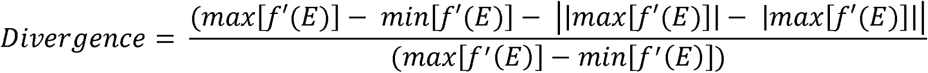

where *f*′(*E*) represents the first derivative of a performance–environment relationship, *f*(*E*), at a particular environmental state (E). Divergence is based on the first derivative of response curves and considers the direction of the performance responses which is crucial for a potential stabilizing effect on aggregate community properties. Furthermore, it produces the largest values for derivatives that are symmetrical around zero. Hence, divergence provided an empirically tractable and intuitive measure of the distribution of species’ fundamental responses.

Note that we used the difference in performance (intrinsic growth rate) from the lower temperature to the higher instead of first derivatives to make divergence applicable to our experimental desig 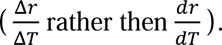. This reflects the fact that microcosms were exposed most of the time to these two temperatures (for simplicity we ignored transitions between temperatures).

We subsequently selected three communities for each richness (3-levels) and environmental treatment (3x3-levels): One with the lowest divergence (often 0, species show the same direction of response, i.e. same sign, unbalanced response distribution), one with the highest divergence (species have different direction and magnitude of responses, balanced responses distribution), and one closest to the mean divergence. Three communities within each of the three richness levels fully-factorially crossed with the three levels of each of the two environmental treatments gives a 3x3x3x3 experimental design. There were 3 replicates of each of the 81 treatment combinations. Thus, the community experiment consisted of 243 experimental units in total. Note that we based this selection purely on divergence and not composition, resulting in some community compositions being used in multiple treatments.

### Sampling plan

The community experiment lasted for sixty days. We sampled every Monday, Wednesday and Friday using video sampling techniques ^36,37^, which yielded time series data of community biomass with 26 time points for each microcosm (6’318 unique samplings). To do so, microcosms were removed from incubators, stirred and a sample was transferred to a counting chamber using a pipette. Subsequently, we recorded 5-s long videos under the microscope and used the R package BEMOVI to process the video footage ^36^. Both video processing and the sampling procedure were performed identically in the response surfaces experiment. Sampling was not performed blind to the conditions of the experiment. However, the actual counting of individuals was performed by automated computer vision ^36^ and thus limited potential for observer effects. We calculated community biomass by adding up the biovolume of individual ciliates for each microcosm and multiplying it with the respective densities, which we approximated as the density of water (1g ^-3^) for all species ^32^. Finally, we quantified temporal stability as the inverse of the coefficient of variation of community biomass for each microcosm, a measure widely used in community experiments ^21,31,32,38^ to quantify temporal stability because it is easily derived from time series data, dimensionless and scale invariant ^22^.

### Measuring imbalance

While divergence predicted temporal stability (Fig. 3, Supplementary Information 1), it has conceptual limitations (Supplementary Information 1, section 5): (1) Divergence only considers the two species with the most extreme responses (the smallest and the largest derivative), ignoring all other members even though they still contribute to overall ecosystem function. (2) If all species in a community respond with the same sign (either exclusively positive or negative first derivatives) divergence will always take the value of zero, meaning that it ignores the magnitude of species responses for those communities. (3) Divergence does not account for the fact that community stability can also result from weak, rather than opposing, responses to environmental change. However, if species respond weakly, it is unlikely to find asynchrony, but one would rather expect population stability to determine stability.

To overcome these limitations, we propose a new and intuitive measure that captures the *imbalance* of species responses. Similarly to divergence, imbalance is based on the idea that differential species responses can compensate for one another at the aggregate ecosystem level, thus stabilising the community through asynchronous dynamics. However, unlike divergence, it quantifies the collective response of the entire community. Imbalance thus quantifies how (im)balanced the species in the system are in their responses to environmental changes. Mathematically, imbalance is the sum of slopes (response sign and strength) of all species within a community:

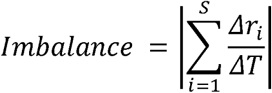

Where

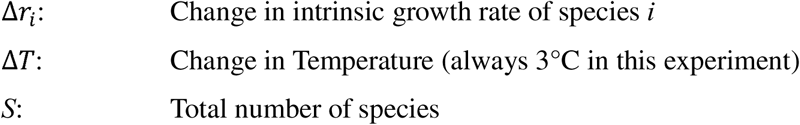

Δ*r*_*i*_ is the change in intrinsic growth rate of species *i* and *S* the total number of species in the community. We take the absolute value of the sum of slopes because we assume a decrease in growth rate of all species in the community to be equally destabilizing as an increase. Values close to zero indicate a balanced community and we would expect high temporal stability.

The larger the value of imbalance, the more variability we expect on the aggregate ecosystem level properties. Again, we used instead of 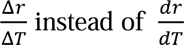 to match our experimental design.

However, under more realistic conditions where communities experience a continuous range of temperatures this would not be appropriate.

Imbalance is a simple summary of species responses that should improve upon the limitations of divergence by considering all species within a community. It gives far from zero values for communities where all species respond with the same sign (exclusively negative/positive slopes). here are two ways by which a small value of imbalance can be achieved: (1) Species respond very differently and compensate one another (both positive and negative slopes). In this case, we expect species to vary asynchronously over time, thus stabilising the community ^10,39^. (2) Species have very weak responses (shallow slopes), meaning that the community is stabilised because populations are stable. As such, we predict that imbalance will outperform existing measures that quantify the distribution of species’ performances because it summarises all species’ responses in a manner that accounts for their effects on both asynchrony and population stability ^16^. For an in-depth comparison of the performance of divergence and imbalance as predictor of temporal stability, please see Supplementary Information 1, section 5.

To further investigate the extent to which the effect of imbalance was influenced by species contributions to ecosystem functioning and other underlying mechanisms (e.g., species interactions), we calculated a realized version of imbalance:

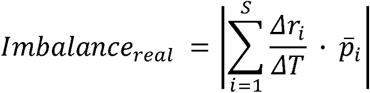

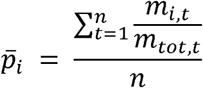

Where:

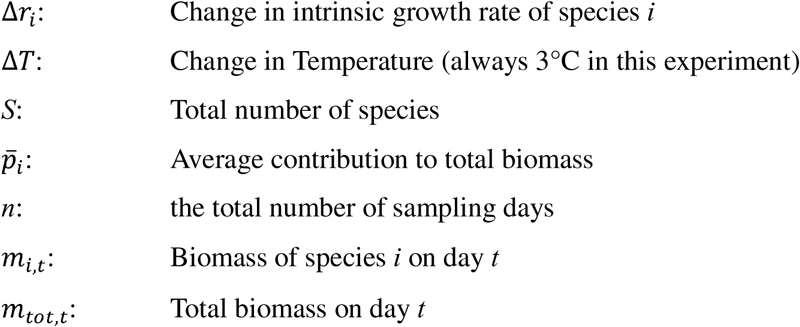

This metric incorporated species-specific contributions by weighting each species’ response according to its proportion of the total community biomass on a given day. The realized imbalance was then averaged across the entire time series to provide a comprehensive measure of its temporal dynamics.

### Analysis

We used a linear mixed model to analyse the results with temporal stability of community biomass as the response variable (measured with the inverse of the coefficient of variation of total community biomass). We log-transformed the response variable to meet model assumptions (i.e., normally distributed residuals). Predictors in our models were: log10(imbalance), nutrients, temperature, and richness. We included an interaction term between nutrients and temperature, and included composition as random effect to account for some compositions being used in more than one treatment combination.

Imbalance is influenced by the slopes of species’ responses across environmental conditions, which are determined by the shape of species response curves and the position of their optima along the environmental gradient. Consequently, we hypothesized that imbalance should depend on the environmental conditions (temperature and nutrients). For example, when environmental fluctuations occur at higher mean temperatures, species’ responses may become more similar (e.g., sharing the same response sign), leading to less symmetrical response distributions and, consequently, higher imbalance (Fig. 7 Supplementary Information 1). This effect can create collinearity among the environment treatments and imbalance.

To disentangle the potential shared explanatory power of imbalance, temperature, and nutrients caused by collinearity we first removed the variance in temperature and nutrients that could be explained by imbalance. This was done by regressing temperature and nutrients on imbalance and extracting the residuals, a method previously used to isolate independent effects among collinear predictors (e.g., ref ^40,41^. The residual temperature and nutrient values thus represented variation independent of imbalance, allowing us to test their effects on stability without confounding influences in the linear mixed model.

The regression models were:

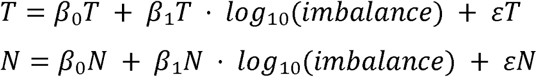

Where:

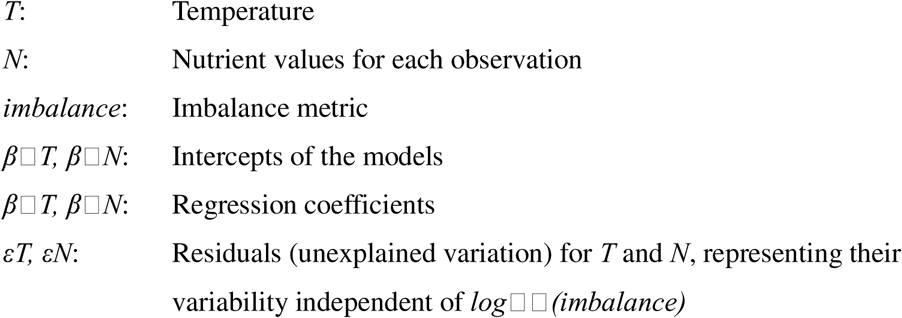

Residuals used in the analysis:

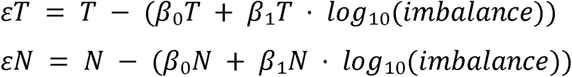

These residuals (ε*T,* ε*N*) were added as new variables (resid_temp and resid_nut) in the dataset.

This approach enabled us to evaluate the relative importance of species richness and imbalance while accounting for environmental variability. By including residual temperature and nutrient values in the model, we ensured that our analysis focused on their independent contributions to stability.

We assessed whether realized and fundamental imbalance differed significantly in their explanatory power by conducting a likelihood ratio test between a model that included both metrics to a reduced model only containing fundamental balance.

To investigate the mechanisms underlying the effect of imbalance on temporal stability, we quantified species asynchrony as proposed by ^21^ using the synchrony function in the ‘codyn’ R package (ref ^42^ version 2.0.5). This measure is calculated by averaging the correlations between each species’ biomass and the total biomass of the rest of the community across all species ^21^. We multiplied (* -1) the values by -1, such that a value of +1 indicates perfect asynchrony, while a value of -1 represents perfect synchrony. For two microcosms, we were not able to calculate asynchrony due to very early extinctions and therefore excluded them from any analysis including asynchrony. We also quantified population stability as the average species level coefficient of variation, weighted by average relative contribution to total community biomass ^16^.

Subsequently, we used structural equation modelling (SEM) as implemented in R by the ‘lavaan’ package (ref ^43^ version 0.6-19) to test our hypothesis that imbalance influences community stability indirectly through its effects on asynchrony and population stability. Due to a certain degree of skew in the distributions of our endogenous variables, we used the Maximum Likelihood estimator with Satorra-Bentler corrections to obtain robust test statistics and standard errors.

## Acknowledgments

This study was founded by the Schweizerischer Nationalfonds zur Förderung der Wissenschaftlichen Forschung project number 320030-231294

