## Supplementary_info1 for "The imbalance of nature: The Role of Species Environmental Responses for Ecosystem Stability"

1 Introduction

2 Load datasets, Data wrangling and Imbalance calculation

3 Biomass

4 Main Results

5 Comparing Divergence and Imbalance

6 Effect RD

7 Linear models

8 Asynchrony

9 Population stability

10 SEM

Code ▾

### Supplementary Materials 1: The balance of nature: Critical Role of Species Environmental Responses for Stability

**Til Hämmig, Francesco Polazzo, Owen L. Petchey, Frank Pennekamp**

**30 January, 2025**

### 1 Introduction

The purpose of this document is to provide a reproducible record of all analyses and figures in the main article. The main article is focused on the effect of response diversity on community stability in fluctuating environments. We are going to look at the effect of the distribution of species responses, richness, temperature and nutrients on community temporal stability. Specifically, we are going to look at the effect of fundamental imbalance (our measurement of the distribution of species' fundamental responses) on temporal stability. Species' fundamental responses to environmental change can stabilise community biomass in two ways: through response diversity and /or through population stability. Thus, as response diversity is thought to stabilize temporal stability of aggregate community properties via asynchrony, we are going to look at the relationship between response diversity and asynchrony. Subsequently, we are going to look at the relationship between population stability and temporal stability of community biomass. Finally, we use a structural equation model to test the direct and indirect effects of balance on temporal stability of community biomass.

In this document, we also analyse the predictive power of imbalance and divergence on temporal stability, and we compare the unique explanatory power of imbalance and divergence. We also look at the interaction between divergence and richness, and between imbalance and richness. Finally, we assess variable importance using the relative importance of predictors in the full model. This part of the document (section 5), is not included in the main article, but it is an important part of the analysis that justify adopting imbalance instead of divergence, which was the original metric used to design the experiment.

This document is produced by an Rmarkdown file that includes code to reproduce from data all results presented in the main article.

### 2 Load datasets, Data wrangling and Imbalance calculation

### 3 Biomass

Let's have a look at the biomass dynamics in the different environmental treatments.

##### 3.0.1 tot biomass plot

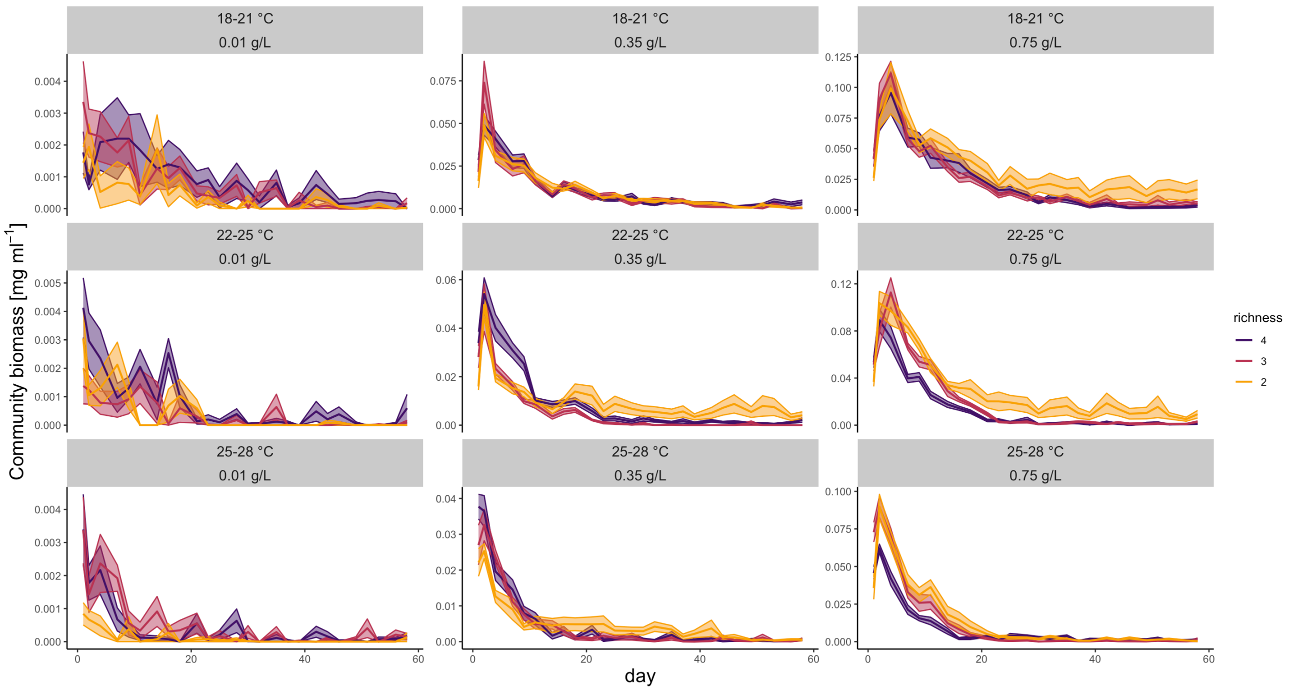


**Figure 1** : Community total biomass during the experiment in different environmental treatments. Different color represent richness levels.

### 4 Main Results

We now look at the main results of the experiment. We are going to look first at the effect of richness, temperature and nutrients on community temporal stability. Then, we are going to look at the relationship between divergence (original response diversity metric) and temporal stability. Finally, we are going to look at the relationship between imbalance and temporal stability.

In the whole analysis, we calculated the temporal stability of total community biomass as the inverse of the coefficient of variation (ICV) (i.e. $\frac{\sigma}{\mu}$).

##### 4.0.1 Effect of T, N and R

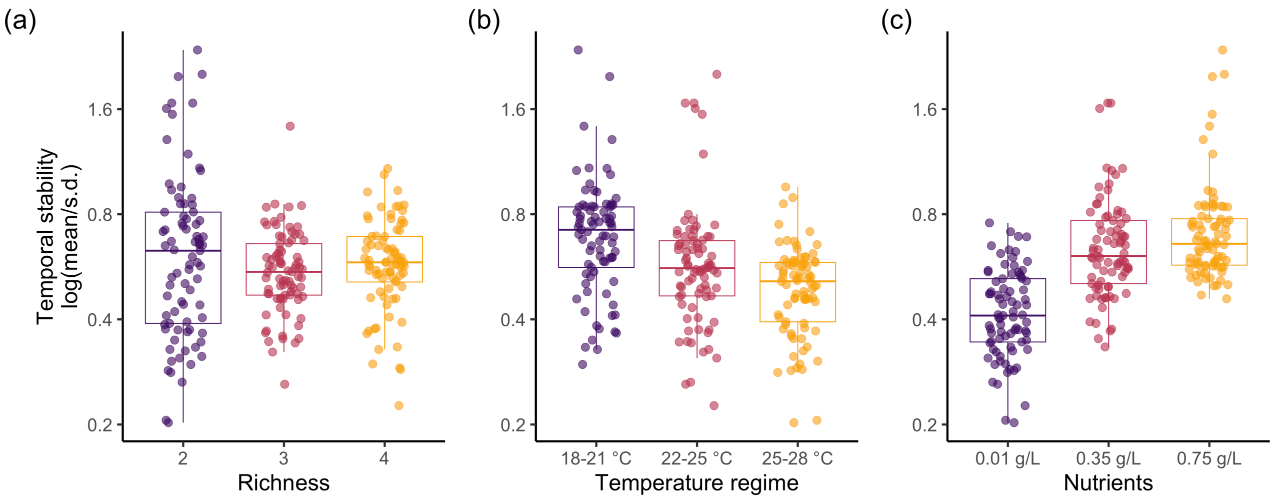


**Figure 2**: Effects of richness (a), temperature (b), and nutrients (c) on community total biomass temporal stability.

We can see that richness does not have a clear effect on community temporal stability, while stability was higher at lower temperature, and nutrients increased community temporal stability.

##### 4.0.2 Effect of Divergence

We look at the relationship between divergence (our original response diversity metric) and stability

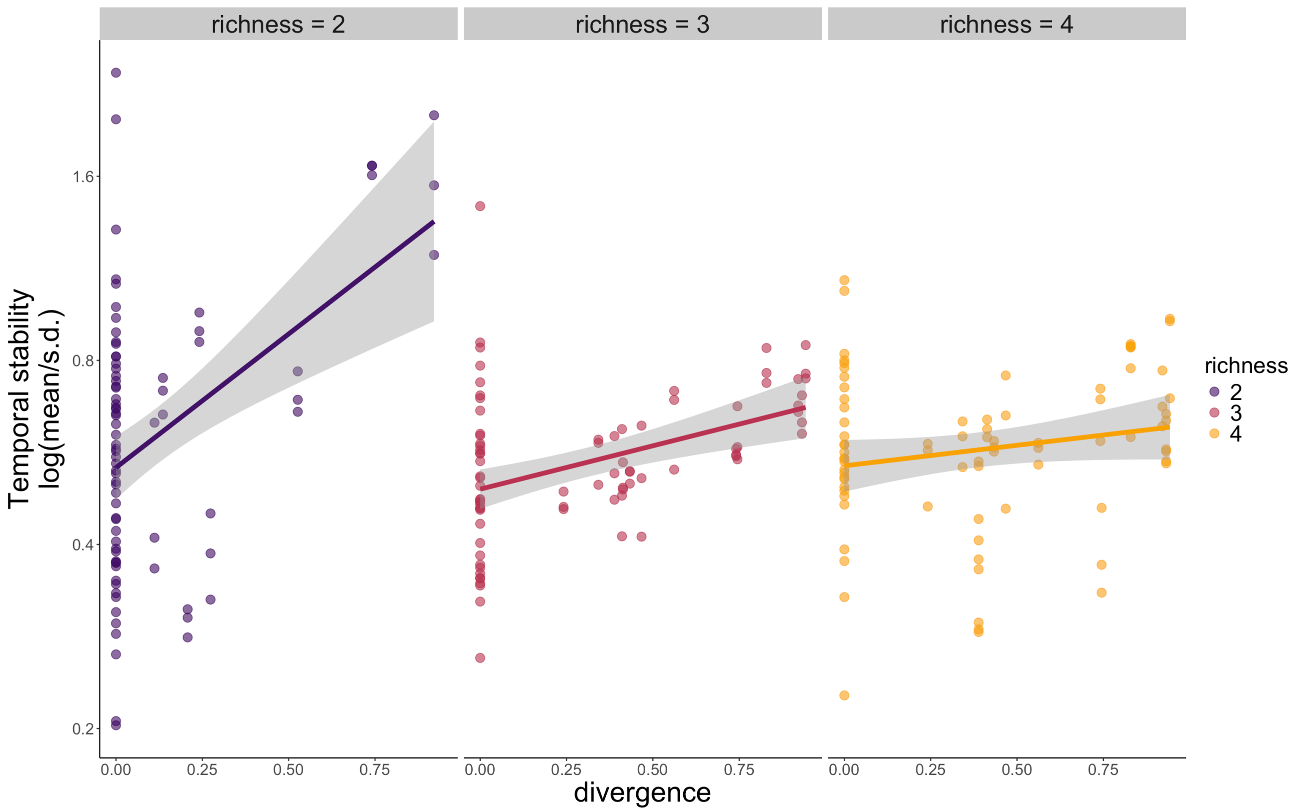


**Figure 3**: Relationship between Divergence and temporal stability of total community biomass.

Divergence is positively related to temporal stability, suggesting that response diversity promotes stability. However, the relationship between divergence and stability becomes weaker as richness increases. We think that this is due to divergence considering only the responses of the 2 most "responding" species. Thus, when species richness increases, disregarding the responses of the other species in the community except the 2 responding the most makes the relationship between response diversity and stability weaker.

This is why, after running the experiment, we developed another metric to measure the distribution of species' responses, which we called **imbalance**, and that is presented in the main text of the publication. Imbalance has several desirable features that make it a more suitable metric than divergence: Independence of richness, higher predictive power, and accounts for the responses of all species in the community (as opposed to divergence that accounts for only the 2 most "responding" species).

Here, we provide extensive evidence of why imbalance is a better metric to measure response diversity than divergence, and thus justifying focusing the analysis around imbalance.

### 5 Comparing Divergence and Imbalance

#### 5.1 Predictive power of Divergence and Imbalance

We first compare how well divergence and imbalance predict stability (predictive power).

##### 5.1.1 Imbalance

Hide

```
#

mod1 <- lm(data=complete_aggr,log10(stability)~log10(balance_f))

# Check model assumptions
#check_model(mod1)
```

##### 5.1.2 Divergence

Hide

```
mod2 <- lm(data=complete_aggr,log10(stability)~(divergence))

# Check model assumptions
#check_model(mod2)
```

**Table 1**: Comparison of model performance of divergence and imbalance as predictors of stability. Model 1 has imbalance as predictor and model 2 has divergence as predictor.

| model | AIC | AICc | BIC | R2 | R2_adjusted | RMSE | Sigma |
|---|---|---|---|---|---|---|---|
| 1 | -89.27328 | -89.17286 | -78.79409 | 0.1917679 | 0.1884142 | 0.1510344 | 0.1516599 |
| 2 | -55.71579 | -55.61538 | -45.23661 | 0.0720796 | 0.0682293 | 0.1618316 | 0.1625017 |

A model with imbalance as predictor performs better than one with divergence as predictor, and it explains more of the variance in stability than divergence.

Moreover, from **Figure 3**, it looks like divergence declines in performance as richness increases. Let's test this analytically. To do than we build a linear model having stability as response variable and either log10(imbalance) or divergence as predictor for each richness level. We then extract the R squared of the models and their *standardised* estimates. (standardized estimates were calculated centering divergence and imbalance using the function scale()).

Show

Show

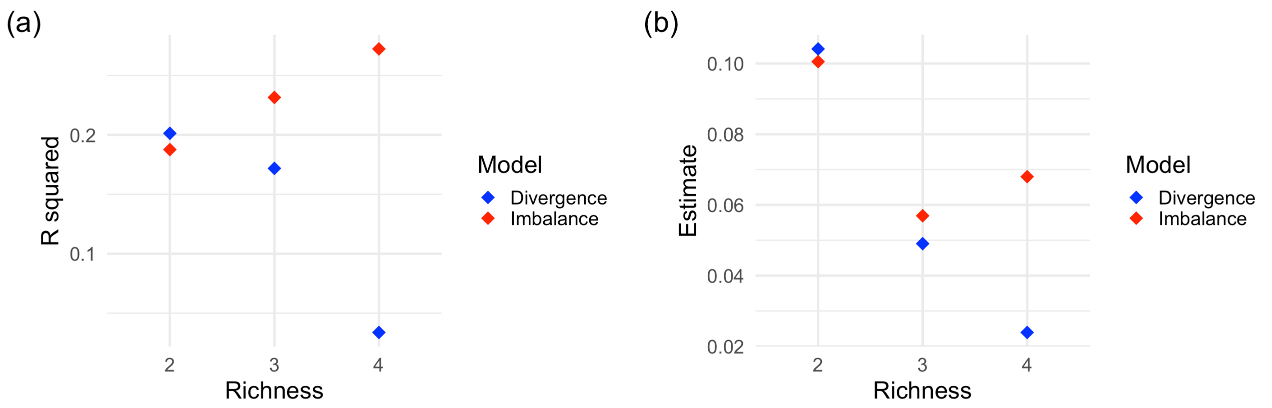


**Figure 4**: Performance comparison of divergence vs imbalance. In (a), the R squared of linear models for divergence and imbalance are shown for each richness level. In (b), the estimates of the linear models for divergence and imbalance are shown for each richness level.

We can see that the R squared of divergence as predictor of stability becomes smaller as richness increases, while the R squared of imbalance as predictor of stability does not (actually increases slightly).

### 5.2 Comparing unique explanatory power of imbalance and divergence

Now we build a linear model were stability is modeled as a function of imbalance and divergence. Then, we compared the variance explained by the full model compared to a model containing either only imbalance or only divergence.

#### 5.2.1 Full model - imbalance and divergence

Hide

```
lm_div_balance <- lm(data=complete_aggr,log10(stability)~log10(balance_f)+diverg
ence)

# Check model assumptions
# check_model(lm_div_balance)
```

#### 5.2.2 model with only divergence

Hide

```
lm_div <- lm(data=complete_aggr,log10(stability)~divergence)

# Check model assumptions
# check_model(lm_div)
```

#### 5.2.3 model with only imbalance

Hide

```
lm_balance <- lm(data=complete_aggr,log10(stability)~log10(balance_f))

# Check model assumptions
# check_model(lm_balance)
```

##### 5.2.4 Comparision full model vs divergence only and imbalance only

**Table 2**: Comparison of model performance of divergence, imbalance and both as predictors of stability. Model 1 has both imbalance and divergence as predictors, model 2 has divergence as predictor, and model 3 has imbalance as predictor.

| model | AIC | AICc | BIC | R2 | R2_adjusted | RMSE | Sigma |
|---|---|---|---|---|---|---|---|
| 1 | -88.97683 | -88.80876 | -75.00458 | 0.1974141 | 0.1907259 | 0.1505060 | 0.1514437 |
| 1 | -55.71579 | -55.61538 | -45.23661 | 0.0720796 | 0.0682293 | 0.1618316 | 0.1625017 |
| 2 | -89.27328 | -89.17286 | -78.79409 | 0.1917679 | 0.1884142 | 0.1510344 | 0.1516599 |

##### 5.2.5 Comparision full model vs imbalance only

**Table 3**: Anova table: a model with both imbalance and divergence as predictors is not significantly different from a model with only imbalance as predictor.

Show

| Term | df.residual | rss | DF | Sum of Squares | F Statistic | p-value |
|---|---|---|---|---|---|---|
| log10(stability) ~ log10(balance_f) + divergence | 240 | 5.504447 | NA | NA | NA | NA |
| log10(stability) ~ log10(balance_f) | 241 | 5.543171 | -1 | -0.03872444 | 1.688429 | 0.195 |

##### 5.2.6 Comparision full model vs imbalance only and divergence only

**Table 4**: Anova table: a model with both imbalance and divergence as predictors is significantly better from a model with only divergence as predictor.

Show

| Term | df.residual | rss | DF | Sum of Squares | F Statistic | p-value |
|---|---|---|---|---|---|---|
| log10(stability) ~ log10(balance_f) + divergence | 240 | 5.504447 | NA | NA | NA | NA |
| log10(stability) ~ divergence | 241 | 6.364040 | -1 | -0.8595933 | 37.47922 | **0.000** |

Overall, imbalance explains more of the variance in stability than divergence, and there is virtually no difference between a model containing only imbalance and the full model.

### 5.3 Interaction divergence and richness

Richness had to be transformed to numeric and to be centered to avoid collinearity with divergence

Hide

```
lm_rich_div <- lm(data=complete_aggr,log10(stability)~divergence*scale(as.numeri
c(richness)))

# check model assumptions
# check_model(lm_rich_div)
```

**Table 5**: Type III anova table of the model with divergence and richness as predictors of stability.

Show

| Term | Sum of Squares | DF | F Statistic | p-value |
|---|---|---|---|---|
| (Intercept) | 11.033652088 | 1 | 442.55571734 | **0.000** |
| divergence | 0.807044347 | 1 | 32.37025122 | **0.000** |
| scale(as.numeric(richness)) | 0.001236238 | 1 | 0.04958505 | 0.824 |
| divergence:scale(as.numeric(richness)) | 0.249582101 | 1 | 10.01064605 | **0.002** |
| Residuals | 5.958668583 | 239 | NA | NA |

Divergence significantly interact with richness, suggesting that the relationship between divergence and stability changes with richness. While an ideal metric of response diversity should be independent of richness.

We repeat the same model using imbalance instead of divergence.

Hide

```
lm_rich_balance <- lm(data=complete_aggr,log10(stability)~log10(balance_f)*scale
(as.numeric(richness)))

# check model assumptions
# check_model(lm_rich_balance)
```

**Table 6**: Type III anova table of the model with imbalance and richness as predictors of stability.

Show

| Term | Sum of Squares | DF | F Statistic | p-value |
|---|---|---|---|---|
| (Intercept) | 9.54928736 | 1 | 414.779284 | **0.000** |
| log10(balance_f) | 1.26534818 | 1 | 54.961191 | **0.000** |
| scale(as.numeric(richness)) | 0.02471247 | 1 | 1.073401 | 0.301 |

| Term | Sum of Squares | DF | F Statistic | p-value |
|---|---|---|---|---|
| log10(balance_f):scale(as.numeric(richness)) | 0.04049552 | 1 | 1.758948 | 0.186 |
| Residuals | 5.50239554 | 239 | NA | NA |

Imbalance does not significantly interact with richness, suggesting that the relationship between imbalance and stability is stable across richness levels.

### 5.4 Variable importance

Finally, we assess variable importance using the relative importance of predictors in the full model. We use the package vip (https://cran.r-project.org/web/packages/vip/vignettes/vip.html (https://cran.r-project.org/web/packages/vip/vignettes/vip.html)) to calculate the relative importance of predictors in the full model. The function vip::vip for multiple linear regression, or linear models (LMs), uses the absolute value of the -statistic as a measure of VI. Motivation for the use of the associated $t$-statistic is given in Bring (1994) [https://www.tandfonline.com/doi/abs/10.1080/00031305.1994.10476059 (https://www.tandfonline.com/doi/abs/10.1080/00031305.1994.10476059)].

Show

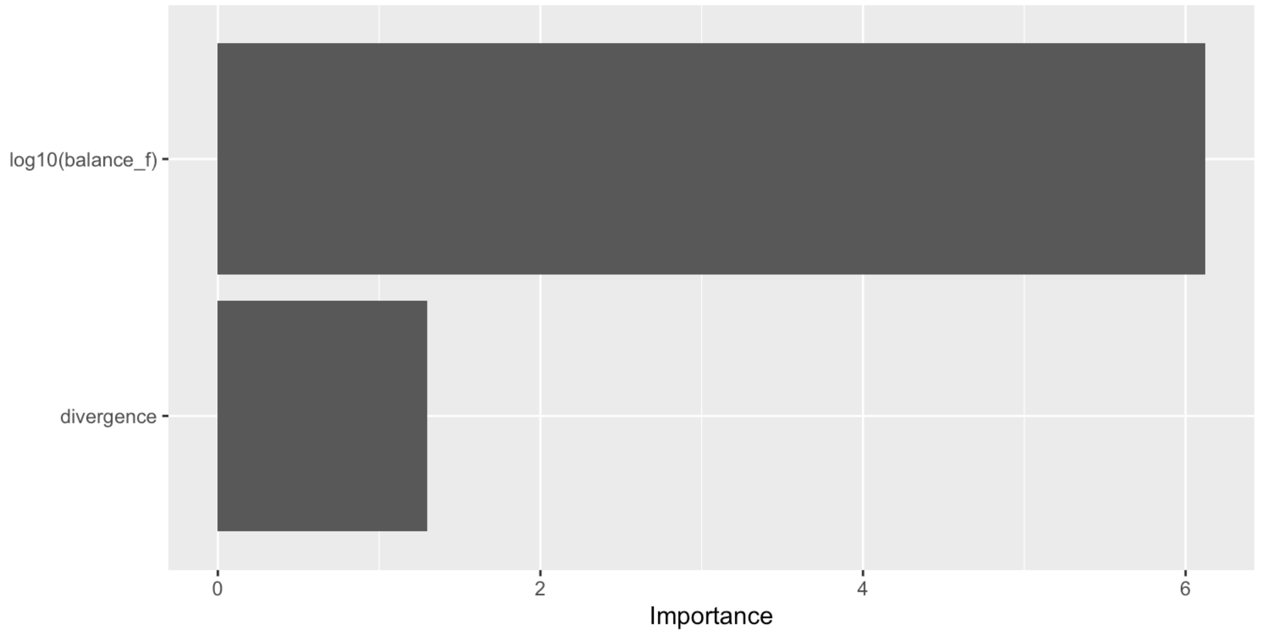


**Figure 5**: Variable importance in the model including both imbalance and divergence as predictors of stability.

We believe that the extensive evidence here provided justifies focusing the analysis around imbalance, and not divergence, as a metric of response diversity. We will thus only look at imbalance for the rest of the analysis.

### 6 Effect RD

We are now going to look at how imbalance affected temporal stability of total community biomass. We are going to look at the relationship between fundamental imbalance (so based only on species response surfaces measured in monoculture), an realised imbalance (measured accounting for species contribution to balance).

This is fundamentally testing our most important hypothesis.

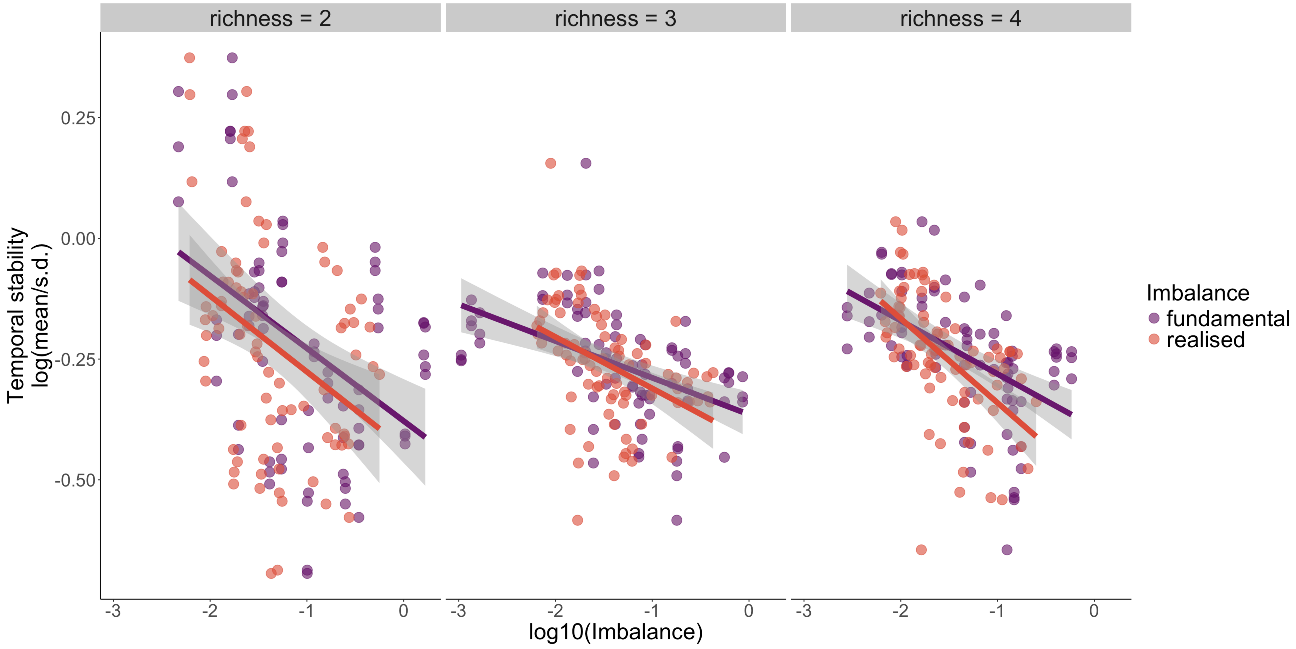


**Figure 6**: Effects of fundamental and realised imbalance on total community biomass temporal stability.

We can see that imbalance is always negatively related to temporal stability, which means that balance in species responses promotes stability across richness levels. Interestingly, we see that there is little difference between fundamental and realised imbalance. Yet, as the richness increases, the relationship between realised imbalance and stability becomes steeper compared to fundamental balance.

But is the difference between fundamental and realised imbalance significant? We can test this using a linear model with both fundamental and realised imbalance as predictors of stability, and one with only fundamental imbalance as predictor of stability, and compare whether the models are significantly different.

### 6.1 Imbalance: realised vs fundamental

Show

**Table 7**: Anova table of the model with only realised balance vs one with both realised and fundamental balance as predictors of stability.

Show

| Term | DF | RSS | df | sumsq | p-value |
|---|---|---|---|---|---|
| log10(stability) ~ log10(balance_f) + scale(nutrients) * scale(temperature) + richness | 236 | 3.624661 | NA | NA | NA |
| log10(stability) ~ log10(balance_f) + log10(balance_r) + scale(nutrients) * scale(temperature) + richness | 235 | 3.609549 | 1 | 0.01511231 | 0.321 |

A model with both fundamental and realised imbalance as predictors improved very little the variance explained by the model. The two models are not significantly different, suggesting that fundamental imbalance captures well the effect of imbalance on stability, and addiing the species contribution to total biomass (realised imbalnce) does not improved the model.

We now compare also the AIC of the two models

Show

```
##              df       AIC
## lm_full_int1  8 -316.2839
## lm_full_int2  9 -315.2992
```

The AIC of the model with only fundamental imbalance is lower than the AIC of the model with both fundamental and realised imbalance, suggesting that the model with only fundamental imbalance is a better model. However, the difference is very small.

### 7 Linear models

#### 7.1 Dependence of Imbalance on Temperature

Since imbalance depends on the slopes of species' responses, and the slopes of species responses between two environmental conditions depend on the shape of species responses and the position of the optimum of the species response curve, we may expect that imbalance depends on temperature. For example, when the environment (i.e. temperature) fluctuates at higher mean temperatures, the species responses may be more similar (i.e. all species have the same sign of response), leading to less symmetrical response distributions, and thus higher imbalance.

We look now at how imbalance changes with temperature, in all possible combinations of species responses for each richness level and each environmental condition. We calculate the sum of slopes of all possible combinations of species responses, and then calculate the imbalance for each combination. We then plot the relationship between imbalance and temperature for each combination of species responses.

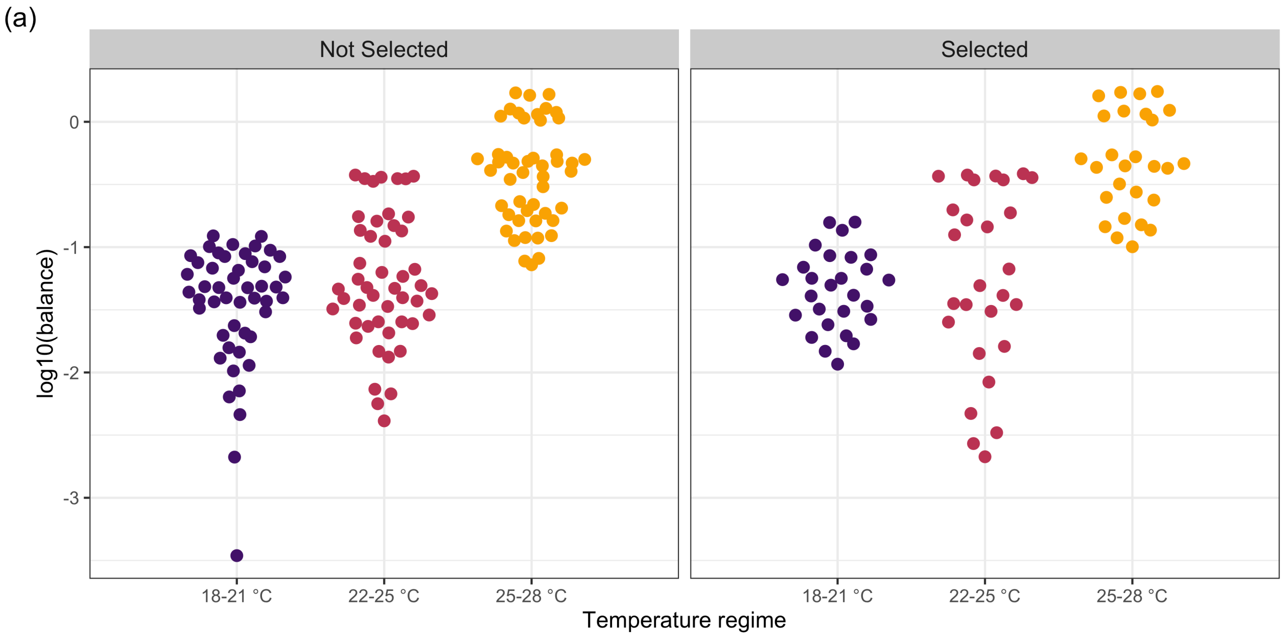


**Figure 7**: Relationship between imbalance and temperature for all possible combinations of species responses. The points represent the imbalance of each combination of species responses The points are colored by temperature regime, and the facets represent whether the combination of species responses was selected for the experiment.

We can see that imbalance increases with temperature, and that the relationship is the same in selected vs non selected community compositions. This suggests that imbalance is intrinsically dependent on temperatures, and that the species responses are more similar when the environment fluctuates at higher temperatures. Inevitably, this will affect the stability of the community, as we have seen in the previous analysis. Yet, it also mean that the effect of imbalance on stability is not independent of temperature, or more generally, of the environmental conditions. Indeed, temperature and nutrients have been used to estimate species' responses, and thus imbalance ineherently depends on the environment.

Let's look at the variance explained by imbalance and the environment on stability.

(a)

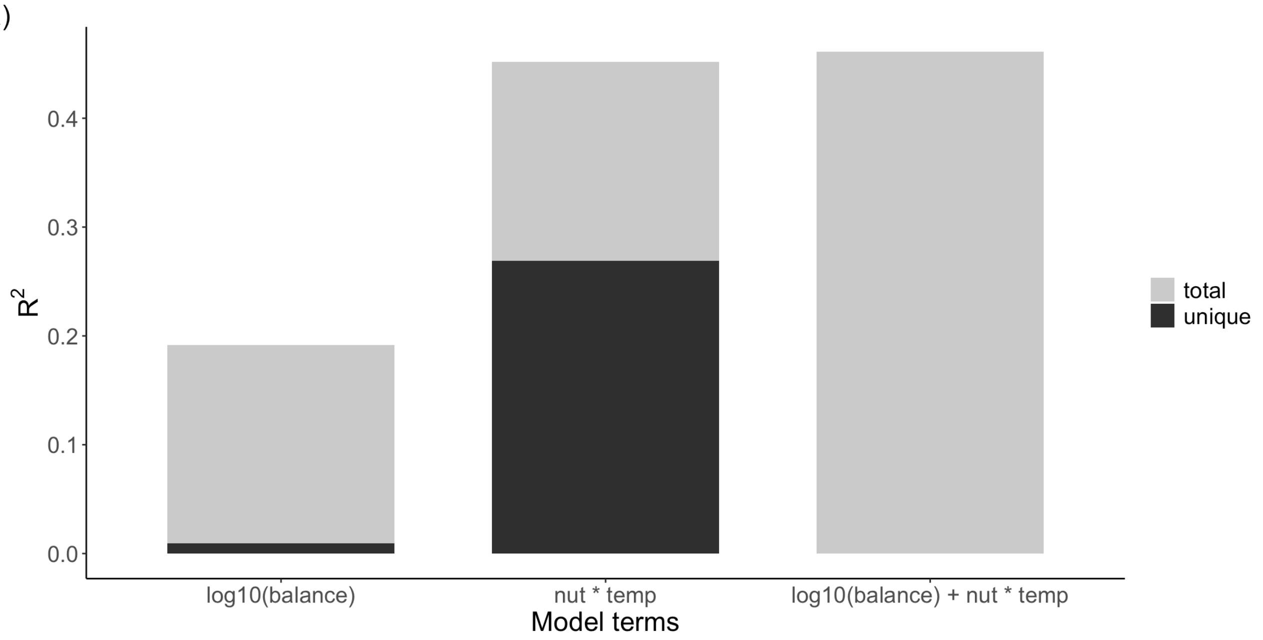


**Figure 8**: Variance explained by imbalance and the environment on stability. The total variance explained by imbalance and the environment is shown in grey, and the unique variance explained by imbalance and the environment is shown in light grey.

95% of the variance explained by balance is shared with nutrients and temperature, which is not suprising considering how imbalance is calculated. However, there is additional variance explained by the environment that is not shared with balance. This suggests that the environment has an independent effect on stability, which is not mediated by imbalance. This may be related to the fact that imbalance was calculated using monoculture data collected at constant temperature, whereas in the community experiment, temperature was fluctuating, and species were interacting.

To disentangle the shared variance between imbalance, temperature, and nutrients, we first removed the variance in temperature and nutrients that could be explained by imbalance. This can be done by regressing temperature and nutrients on imbalance and extracting the residuals, a method previously used to isolate independent effects among collinear predictors (e.g., Dormann et al. 2013 (https://nsojournals.onlinelibrary.wiley.com/doi/full/10.1111/j.1600-0587.2012.07348.x); Graham 2003 (https://esajournals.onlinelibrary.wiley.com/doi/full/10.1890/02-3114?casa_token=tjLz0BPcLf4AAAAA%3AYjunSb9xIIaHNsO0YLwGDjWzZswdTkVOWfcsBfIC3WjrA-n5B_94cOP2LFP5WIXMhunDwZOPFXCQqg)). The residual temperature and nutrient values thus represented variation independent of imbalance, allowing us to test their effects on stability without confounding influences of shared variance with imbalance. This approach enabled us to evaluate the relative importance of species richness and imbalance while accounting for environmental variability. By including residual temperature and nutrient values in the model, we ensured that our analysis focused on their independent contributions to stability.

### 7.2 Control effect of environment on balance

Now, we can build a linear model with imbalance, richness, the residuals of temperature and nutrients and their interaction as predictors of stability. We also use composition as random effect to account for some compositions being used in more than one treatment combination.

We also want to test whether the inclusion of random slopes for log10(balance_f) improves the model's ability to explain variability in log10(stability) among compositions. We will compare the model with random slopes for log10(balance_f) to the model without random slopes for log10(balance_f) using a Likelihood Ratio Test.

```
## Data: complete_aggr_2
## Models:
## model_no_random: log10(stability) ~ log10(balance_f) + richness + resid_temp
* resid_nut + (1 | composition)
## model_with_random: log10(stability) ~ log10(balance_f) + resid_temp * resid_n
ut + richness + (1 + log10(balance_f) | composition)
##                   npar     AIC     BIC logLik deviance  Chisq Df Pr(>Chisq)
## model_no_random      9 -501.11 -469.68 259.56  -519.11
## model_with_random   11 -499.25 -460.83 260.63  -521.25 2.1398  2      0.343
```

A model with random slopes for log10(balance_f) is not better than one without. Thus, we will use the model without random slopes for log10(balance_f) for the rest of the analysis.

Show

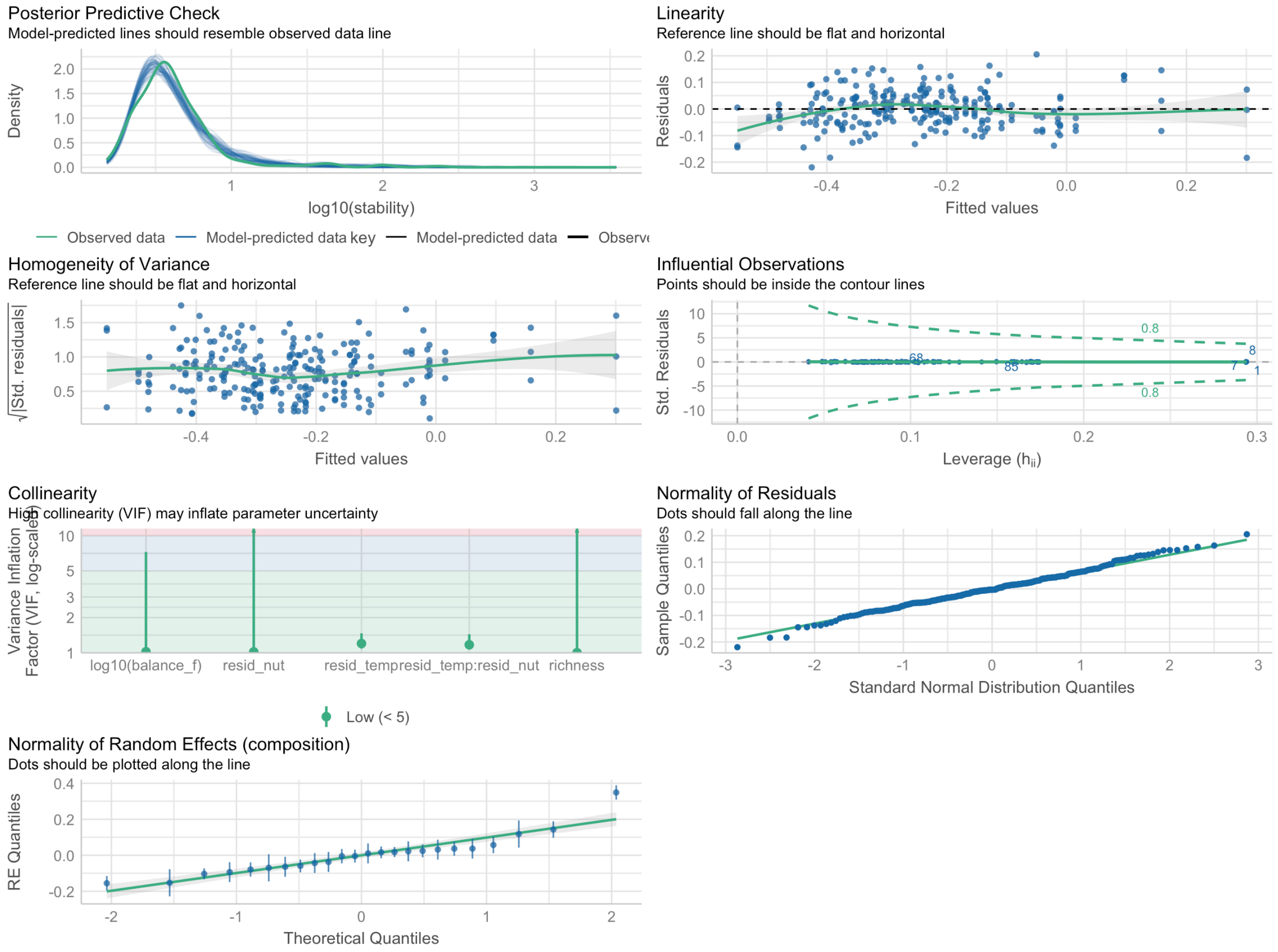


(#fig:model_check_int)model check 1.

**Figure 9**: Model check for the linear mixed-effects model with balance, richness, and the residuals of temperature and nutrients as predictors of stability. The model fits well the data.

**Table 10**: Linear mixed-effects model results for the effects of balance, richness, and the residuals of temperature and nutrients on community stability. Estimates are presented with 95% confidence intervals and p-values.

| Linear Regression Results | | | | |
|---|---|---|---|---|
| **Predictor** | **Estimate** | **95% CI**[1] | **t-value** | **p-value** |
| **Intercept** | -0.32 | -0.39, -0.24 | -8.84 | **<0.001** |
| **log10(balance_f)** | -0.08 | -0.09, -0.06 | -9.95 | **<0.001** |
| **richness** | | | | |
| richness3 - richness2 | -0.01 | -0.15, 0.12 | -0.27 | >0.9 |
| richness4 - richness2 | 0.01 | -0.15, 0.17 | 0.13 | >0.9 |
| richness4 - richness3 | 0.02 | -0.14, 0.18 | 0.35 | >0.9 |
| **resid_temp** | -0.10 | -0.12, -0.08 | -10.55 | **<0.001** |
| **resid_nut** | 0.25 | 0.21, 0.28 | 13.80 | **<0.001** |
| **resid_temp * resid_nut** | -0.09 | -0.15, -0.03 | -2.93 | **0.004** |

[1] CI = Confidence Interval

The linear mixed-effects model was fitted by maximum likelihood (ML) with log-transformed stability as the response variable. The fixed effects included log10(balance_f), richness, the residuals of temperature (resid_temp), and nutrients (resid_nut), as well as their interaction term. The random effect was specified for the composition group, which accounted for potential variability between different composition categories.

The model's results indicated that log10(balance_f) (Estimate = -0.078, SE = 0.0079, p < 2e-16), resid_temp (Estimate = -0.099, SE = 0.0093, p < 2e-16), and resid_nut (Estimate = 0.248, SE = 0.018, p < 2e-16) all had significant effects on stability, with both resid_temp and resid_nut showing large, statistically significant relationships with stability. Additionally, the interaction term resid_temp:resid_nut (Estimate = -0.091, SE = 0.031, p = 0.0037) was also significant, indicating that the combined effects of temperature and nutrients on stability are not independent, but interact in shaping stability.

Richness levels (richness3 and richness4) did not show significant effects on stability (p > 0.77), suggesting that, within the context of this model, variations in richness do not explain substantial variation in community stability.

**Table 11**: ANOVA table of the linear mixed-effects model with balance, richness, and the residuals of temperature and nutrients as predictors of stability.

| Term | F Statistic | DF | p-value |
|---|---|---|---|
| (Intercept) | 78.2322254 | 1 | **0.000** |
| log10(balance_f) | 99.0517826 | 1 | **0.000** |
| richness | 0.1612692 | 2 | 0.923 |
| resid_temp | 111.3786982 | 1 | **0.000** |

| Term | F Statistic | DF | p-value |
|---|---|---|---|
| resid_nut | 190.4777948 | 1 | **0.000** |
| resid_temp:resid_nut | 8.5784507 | 1 | **0.003** |

### 8 Asynchrony

Response diversity (one of the stabilisng effects captured by imbalance) has been suggested as a mechanism that promotes temporal stability of community biomass by promoting species asynchrony.

We thus calculated the asynchrony index suggested by Gross et al. 2014 (https://www.journals.uchicago.edu/doi/epdf/10.1086/673915) to calculate the effect of asynchrony on temporal stability and to see how response diversity relate to asynchrony. The index ranges between -1 and 1, with -1 indicating perfect asynchrony and 1 being perfectly synchronous, and 0 indicating random variation.

##### 8.0.1 Plot stability vs. Asynchrony Gross

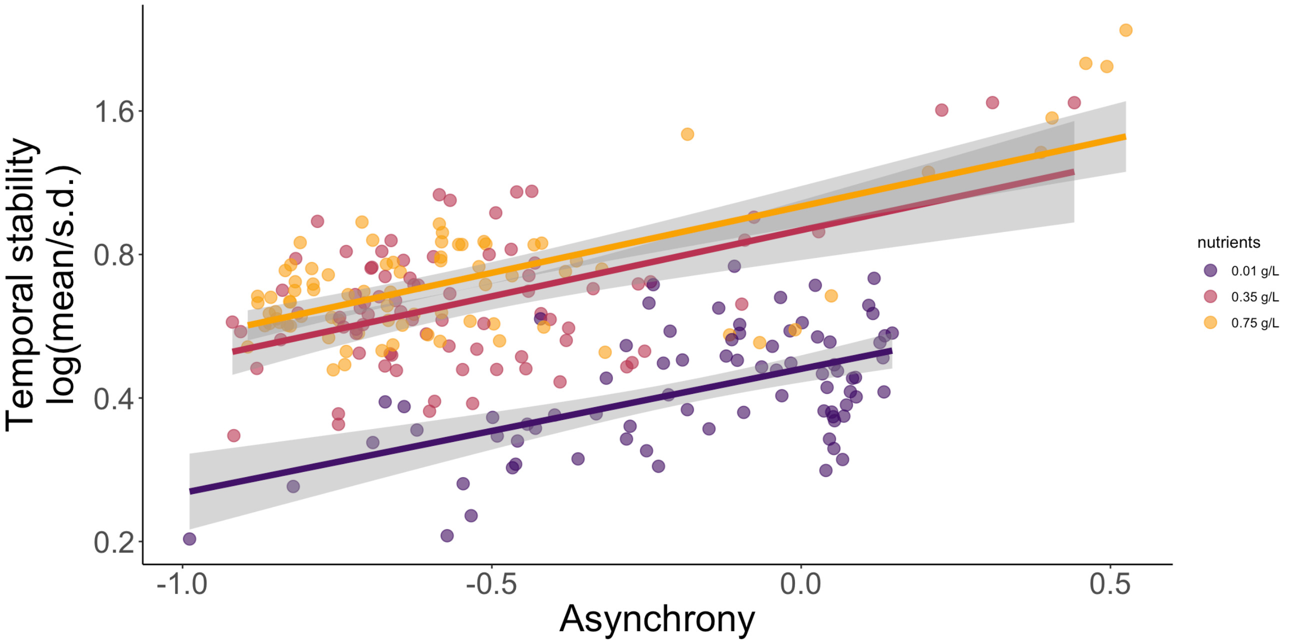


**Figure 10**: Relationship between temporal stability and asynchrony (Gross) divided by nutrient level.

##### 8.0.2 Plot Asynchrony Gross vs fundamental imbalance

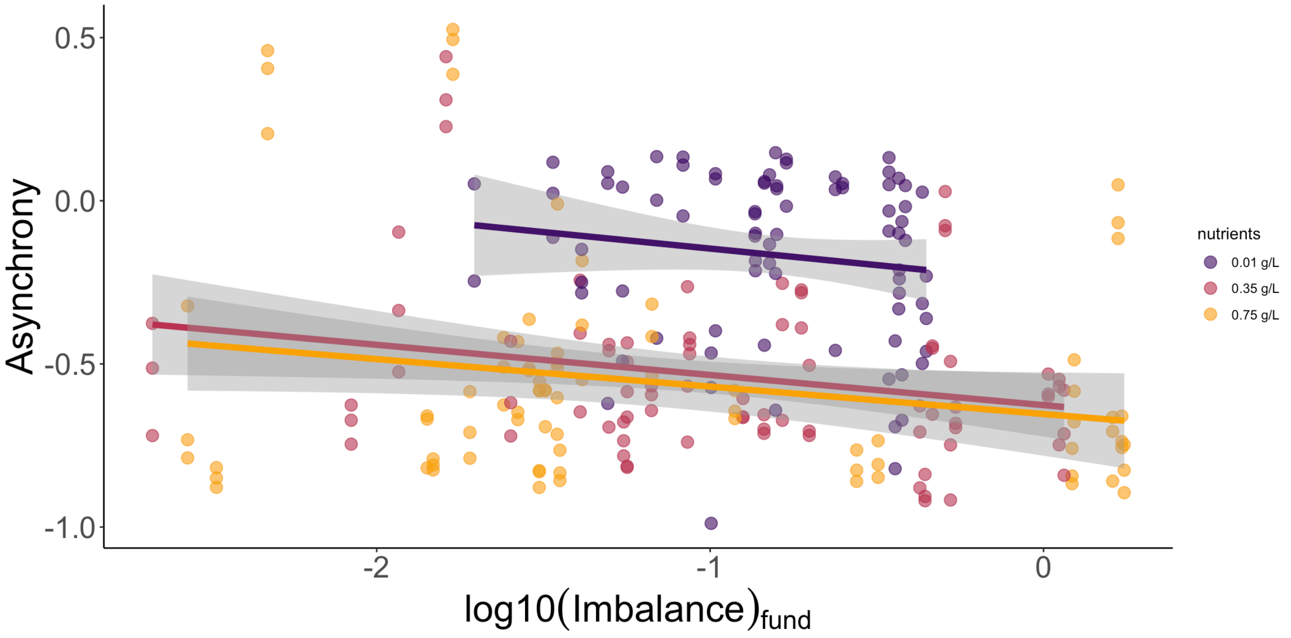


**Figure 11**: Relationship between asynchrony (Gross) and fundamental imbalance divided by nutrient level.

#### 8.1 Eveness

Evenness in species biomass has been identified as an important factor potentially influencing ecosystem stability Thibaut & Connolly 2013 (https://onlinelibrary.wiley.com/doi/full/10.1111/ele.12019). In the context of our experiment, evenness in species biomass could help explaining why there is little difference between fundamental and realized imbalance. If evenness is high, then all species contribute similarly to total biomass. In this case, weighting for species-biomass contribution to total biomass (realized), should not fundamentally change the result, compared to an un-weighted (fundamental) measurement.

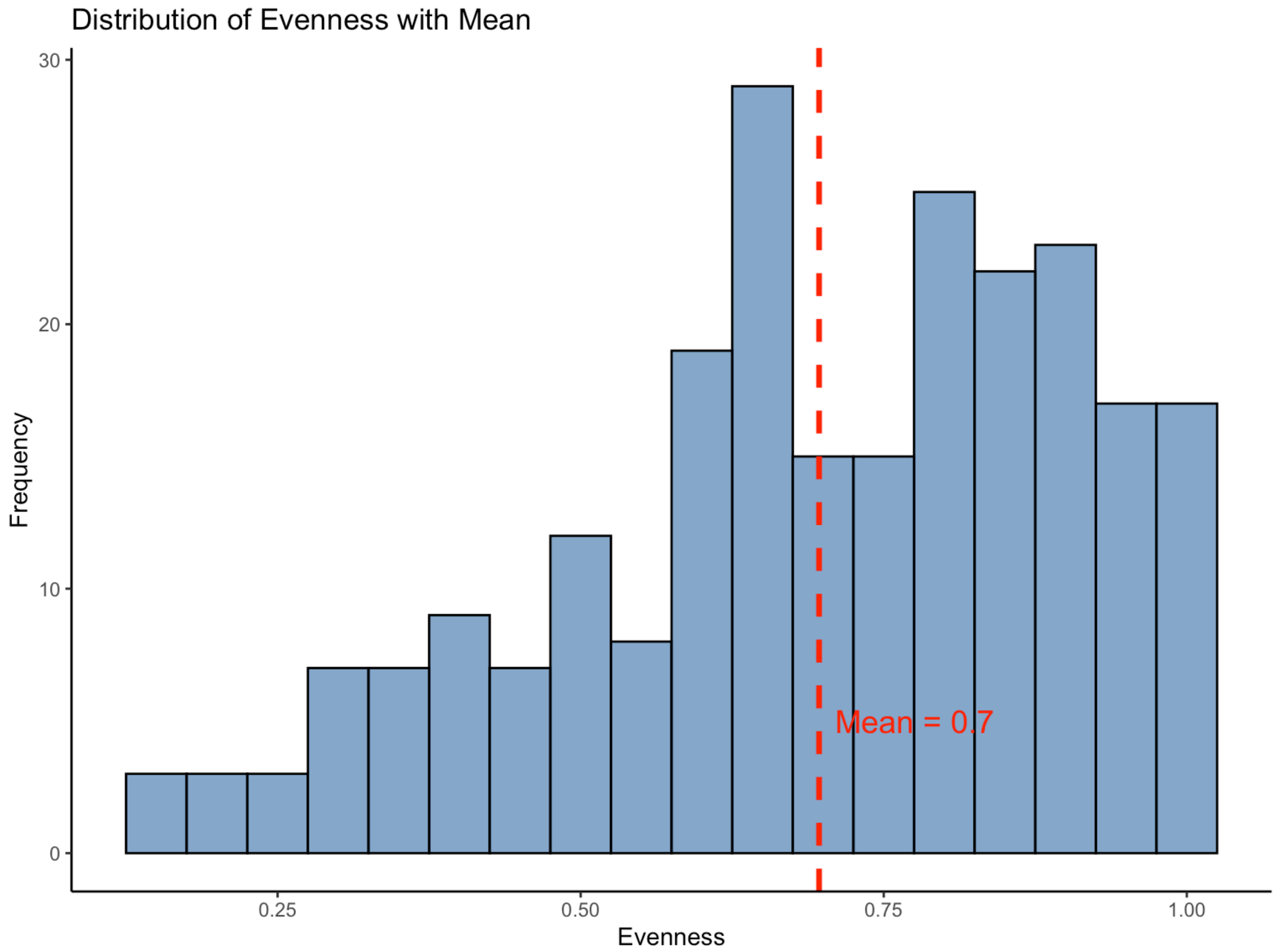


**Figure 12**: Distribution of species evenness across experimental communities. The histogram represents the frequency of observed evenness values, while the red dashed line indicates the mean evenness (0.7). This highlights the central tendency of evenness across the dataset and its variation among communities.

Evenness was indeed generally high in our experimental communities, suggesting another potential factor reducing the potential difference between fundamental and realized balance.

### 9 Population stability

The relationship between community stability and the stability of the individual populations that make up the community is a key question in ecology. Importantly, ecosystem stability can result from low population stability, if populations fluctuate asynchronously, or from high population stability, if populations do not fluctuate much. Synthesis of the literature suggests diversity can have a positive or negantive effect on population stability Campbell et al 2010 (https://nsojournals.onlinelibrary.wiley.com/doi/full/10.1111/j.1600-0706.2010.18768.x) and (Xu et al 2021)[https://onlinelibrary.wiley.com/doi/full/10.1111/ele.13777 (https://onlinelibrary.wiley.com/doi/full/10.1111/ele.13777)].

Theoretical work has suggested that community stability is a product of two quantities: the (a) synchrony of population fluctuations, and an average species-level population stability that is weighted by relative abundance Thibaut & Connolly 2013 (https://onlinelibrary.wiley.com/doi/full/10.1111/ele.12019).

Critically, a imbalance value close to zero can result from high response diversity, but also from high population stability (population biomass does not change largely over time). We want to look now at whether our new metric of imbalance can capture these two stabilising mechanisms.

Thus, we calculate species-level population stability weighted by relative abundance and look at how it relates to ecosystem stability.

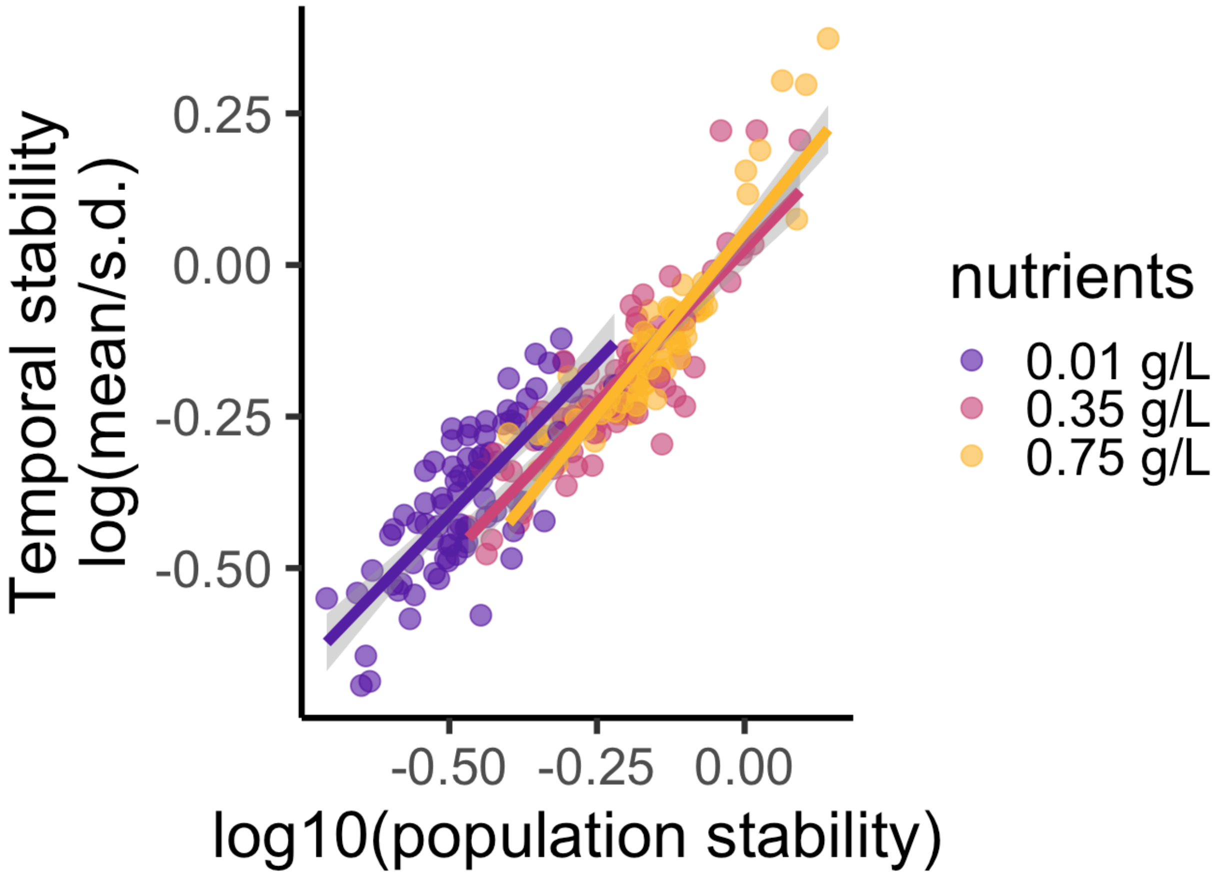


**Figure 13**: Relationship between log10 of population stability and log 10 of ecosystem stability.

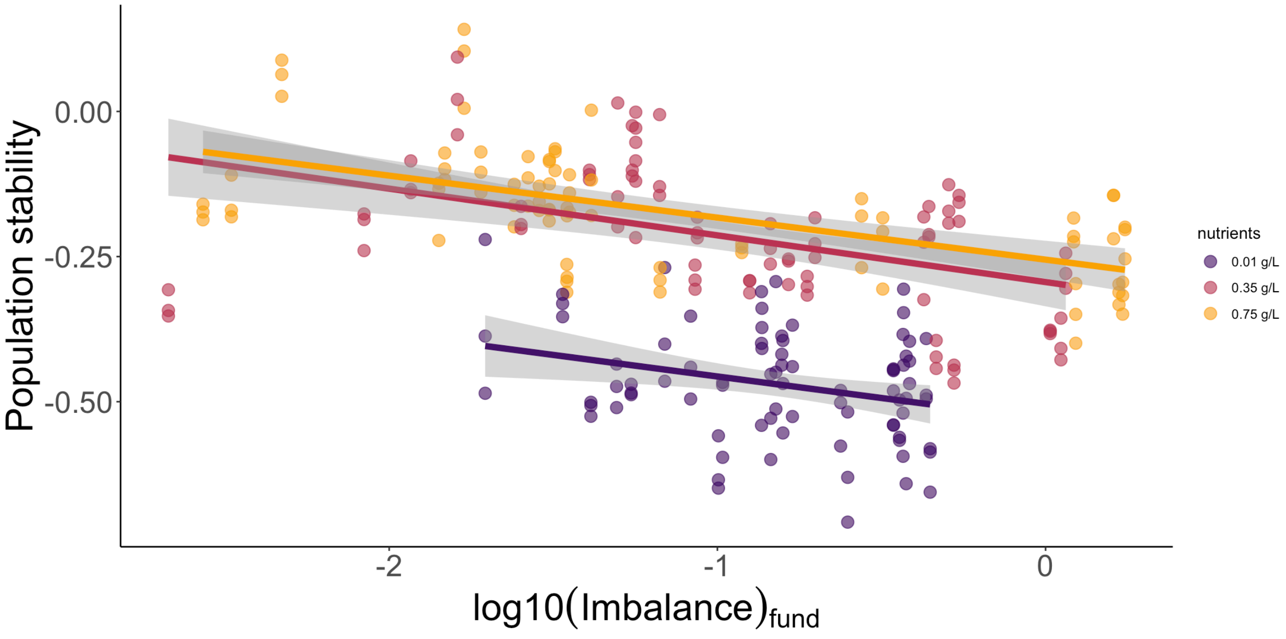


**Figure 14**: Relationship between fundamental imbalance and population stability divided by nutrient level.

### 10 SEM

Finally, we use a structural equation model (SEM) to explore how stability is influenced by asynchrony, population stability, imbalance and, nutrient levels. In order to develop a hypothesis regarding the influence of stability, we have drawn on existing literature. This has enabled us to posit that stability is influenced by two key factors: asynchrony and population stability. In turn, these are influenced by balance and, in our particular case, by nutrient levels.

```
## lavaan 0.6-19 ended normally after 1 iteration
##
##   Estimator                                         ML
##   Optimization method                           NLMINB
##   Number of model parameters                        12
##
##   Number of observations                           241
##
## Model Test User Model:
##                                               Standard      Scaled
##   Test Statistic                                 2.502       2.362
##   Degrees of freedom                                 3           3
##   P-value (Chi-square)                           0.475       0.501
##   Scaling correction factor                                  1.059
##     Satorra-Bentler correction
##
## Model Test Baseline Model:
##
##   Test statistic                               852.984     819.012
##   Degrees of freedom                                 9           9
##   P-value                                        0.000       0.000
##   Scaling correction factor                                  1.041
##
## User Model versus Baseline Model:
##
##   Comparative Fit Index (CFI)                    1.000       1.000
##   Tucker-Lewis Index (TLI)                       1.002       1.002
##
##   Robust Comparative Fit Index (CFI)                         1.000
##   Robust Tucker-Lewis Index (TLI)                            1.002
##
## Loglikelihood and Information Criteria:
##
##   Loglikelihood user model (H0)                516.122     516.122
##   Loglikelihood unrestricted model (H1)             NA          NA
##
##   Akaike (AIC)                               -1008.245   -1008.245
##   Bayesian (BIC)                              -966.427    -966.427
##   Sample-size adjusted Bayesian (SABIC)      -1004.464   -1004.464
##
## Root Mean Square Error of Approximation:
##
##   RMSEA                                          0.000       0.000
##   90 Percent confidence interval - lower         0.000       0.000
##   90 Percent confidence interval - upper         0.101       0.097
##   P-value H_0: RMSEA <= 0.050                    0.692       0.723
##   P-value H_0: RMSEA >= 0.080                    0.127       0.107
##
##   Robust RMSEA                                               0.000
##   90 Percent confidence interval - lower                     0.000
##   90 Percent confidence interval - upper                     0.102
##   P-value H_0: Robust RMSEA <= 0.050                         0.703
##   P-value H_0: Robust RMSEA >= 0.080                         0.126
##
## Standardized Root Mean Square Residual:
```

```
##
##   SRMR                                           0.018       0.018
##
## Parameter Estimates:
##
##   Standard errors                             Robust.sem
##   Information                                   Expected
##   Information saturated (h1) model            Structured
##
## Regressions:
##                      Estimate  Std.Err  z-value  P(>|z|)   Std.lv  Std.all
##   stability ~
##     asynchrny_Grss      0.157    0.011   14.493    0.000    0.157    0.326
##     pop_stability       0.957    0.022   42.596    0.000    0.957    0.987
##   asynchrony_Gross ~
##     log_balance_f      -0.093    0.034   -2.767    0.006   -0.093   -0.185
##     nutrients          -0.211    0.023   -9.072    0.000   -0.211   -0.503
##   pop_stability ~
##     log_balance_f      -0.073    0.011   -6.852    0.000   -0.073   -0.292
##     nutrients           0.136    0.008   17.854    0.000    0.136    0.653
##
## Intercepts:
##                    Estimate  Std.Err  z-value  P(>|z|)   Std.lv  Std.all
##    .stability         0.098    0.011    8.847    0.000    0.098    0.594
##    .asynchrny_Grss   -0.090    0.055   -1.625    0.104   -0.090   -0.262
##    .pop_stability    -0.630    0.018  -34.512    0.000   -0.630   -3.702
##
## Variances:
##                    Estimate  Std.Err  z-value  P(>|z|)   Std.lv  Std.all
##    .stability         0.003    0.000    7.632    0.000    0.003    0.099
##    .asynchrny_Grss    0.088    0.011    8.214    0.000    0.088    0.753
##    .pop_stability     0.012    0.001   11.568    0.000    0.012    0.406
##
## R-Square:
##                    Estimate
##     stability         0.901
##     asynchrny_Grss    0.247
##     pop_stability     0.594
```

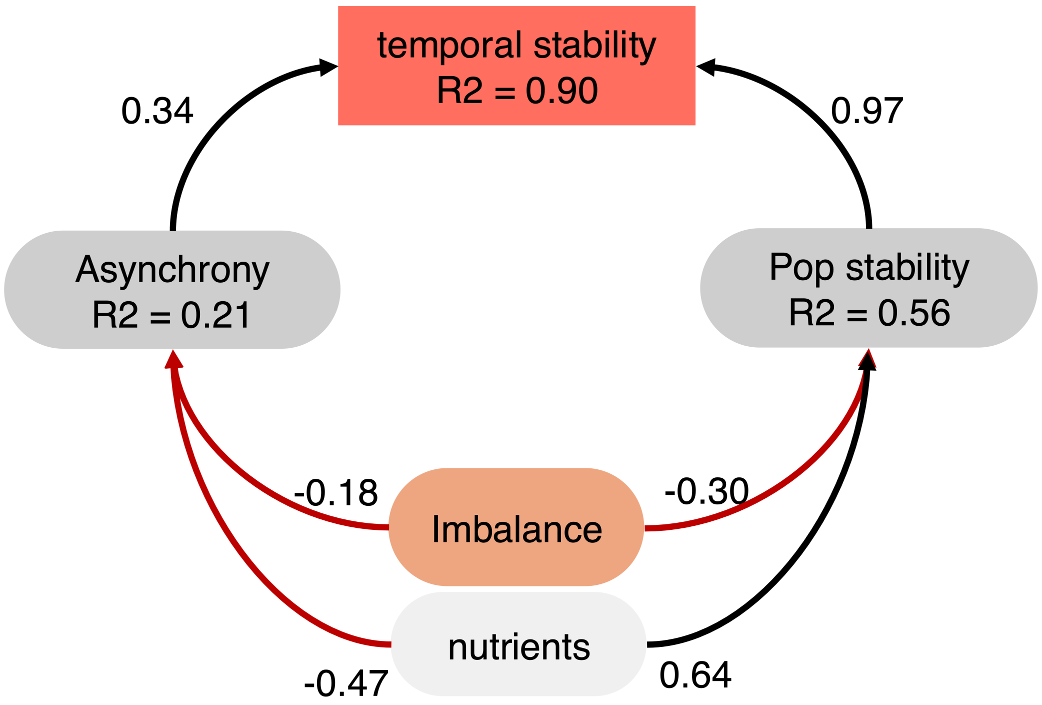


$\chi 2 = 1.62, df = 3, p - value = 0.69, CFI = 1.0, RMSEA = 0.0$

Figure 10.1: SEM.

**Figure 15**: Structural equation model (SEM) showing the relationships between stability, asynchrony, population stability, imbalance, and nutrient levels. The model includes standardized regression coefficients and fit indices.

**Model Fit Indices** The model fit indices suggest that the model fits the data well.

*Chi-Square Test (User Model)*: The chi-square test statistic for the user model is χ 2 =1.626 (scaled = 1.465) with 3 degrees of freedom and a p-value of 0.653 (scaled = 0.690). This indicates a good fit, as the test is non-significant, suggesting no significant difference between the observed and model-implied covariance matrices.

*Comparative Fit Index (CFI)* and *Tucker-Lewis Index (TLI)*: Both CFI and TLI values are 1.000, indicating an excellent model fit. Values close to or above 0.95 are generally considered good.

*Root Mean Square Error of Approximation (RMSEA)*: The RMSEA is 0.000, with a 90% confidence interval ranging from 0 to 0.090 (scaled = 0.080). This indicates a very good fit, as RMSEA values below 0.05 are ideal, and values below 0.08 are acceptable. The p-values for the RMSEA hypothesis tests suggest strong support for a close fit (RMSEA <= 0.05) and little evidence for a poor fit (RMSEA >= 0.08).

*Standardized Root Mean Square Residual (SRMR)*: The SRMR value is 0.017, which is also within the acceptable range (values below 0.08 are generally considered good). Overall, the fit indices suggest that the model is an excellent fit for the data.

**Regression Paths and Interpretation**

**Stability Regressions**

*Stability ~ Asynchrony_Gross (asynchrny_Grss)*: The standardized estimate for the effect of asynchrony on stability is 0.340 (p < 0.001), indicating a significant positive association. Higher asynchrony in species dynamics is associated with increased community stability.

*Stability ~ Population Stability (pop_stability)*: The standardized estimate is 0.977 (p < 0.001), showing a strong positive relationship. This suggests that community stability is highly dependent on the stability of individual populations within the community.

**Asynchrony_Gross Regressions**

*Asynchrony_Gross ~ Log10(Balance):* The standardized estimate is -0.176 ($p = 0.013$), indicating a significant negative effect. Higher imbalance leads to lower asynchrony, suggesting that as imbalance increases, species within the community fluctuate more synchronously.

*Asynchrony_Gross ~ Nutrients:* The standardized estimate is -0.469 ($p < 0.001$), showing a strong negative relationship. Higher nutrient levels appear to reduce asynchrony, possibly by causing similar responses across species.

**Population Stability Regressions**

*Population Stability ~ Log10(Balance)*: The standardized estimate is -0.296 ($p < 0.001$), indicating that higher imbalance is associated with lower population stability.

*Population Stability ~ Nutrients*: The standardized estimate is 0.635 ($p < 0.001$), showing that higher nutrient levels are associated with increased population stability, likely because nutrients enhance conditions that support stable population dynamics.

**Variances and R-Squared Values** *R-Squared for Stability*: The model explains 90.4% of the variance in community stability, indicating strong predictive power.

*R-Squared for Asynchrony_Gross*: The model explains 21.9% of the variance in asynchrony, which is moderate.

*R-Squared for Population Stability*: The model explains 56.2% of the variance in population stability, showing that nutrients and balance are important but not the only factors influencing it.

*Summary Interpretation* Model Fit: The model has an excellent fit, as indicated by the fit indices. Stability: Community stability is strongly influenced by both population stability and asynchrony among species, with population stability being the stronger predictor. Asynchrony and Imbalance: Asynchrony decreases with increasing imbalance and nutrients, suggesting that these factors promote more synchronized fluctuations among species. Population Stability and Nutrients: Higher nutrient levels are associated with increased population stability, suggesting that nutrient availability supports stable population dynamics. Conversely, higher imbalance is associated with decreased population stability.
