## Supplementary_info2 for "The imbalance of nature: The Role of Species Environmental Responses for Ecosystem Stability"

1 Intention

2 Response diversity with known direction of environmental change

2.1 Temperature fluctuations

2.2 Get the slopes (change in species performance from Temperature 1 to Temperature 2 and vice versa)

2.3 Calculate diveregence for all possible compositions

2.4 Calculate diveregence

Code ▾

### Supplementary materials for: The balance of nature: Critical Role of Species Environmental Responses for Stability

Francesco Polazzo, Til Hämmig, Owen L. Petchey, Frank Pennekamp

23 January, 2025

### 1 Intention

The goal of this document is to calculate the growth rate (r) and the carrying capacity (K) of 6 species of ciliates that we grew individually in a replicated (n = 3) factorial experiment with 5 temperatures (18, 21, 24, 26, and 28), 5 nutrients levels and their combinations (= 25 treatments) for 3 weeks. The calculated r and K are then going to be used as the traits to calculate the response diversity of communities composed of 2, 3, and 4 species to all possible changes in temperature and nutrients. The calculated response diversity will inform us on which communities will be used in the following experimental step. We are going to calculate the growth rate as the slope of the regression of ln(Nt) where: ln is natural log and Nt is the population density at time t, during the period of exponential growth. We are going to set initially the period of exponential growth as the first 6 days, but we are going to visually check whether this choice is correct. K will be calculated as the highest population biomass for each population during the experiment.

Let's start loading the data set and creating a subset for calculating r The data set loaded here is a reanalysis of monoculture data, where different bemovi settings have been used to stay consistent between poly and monoculture

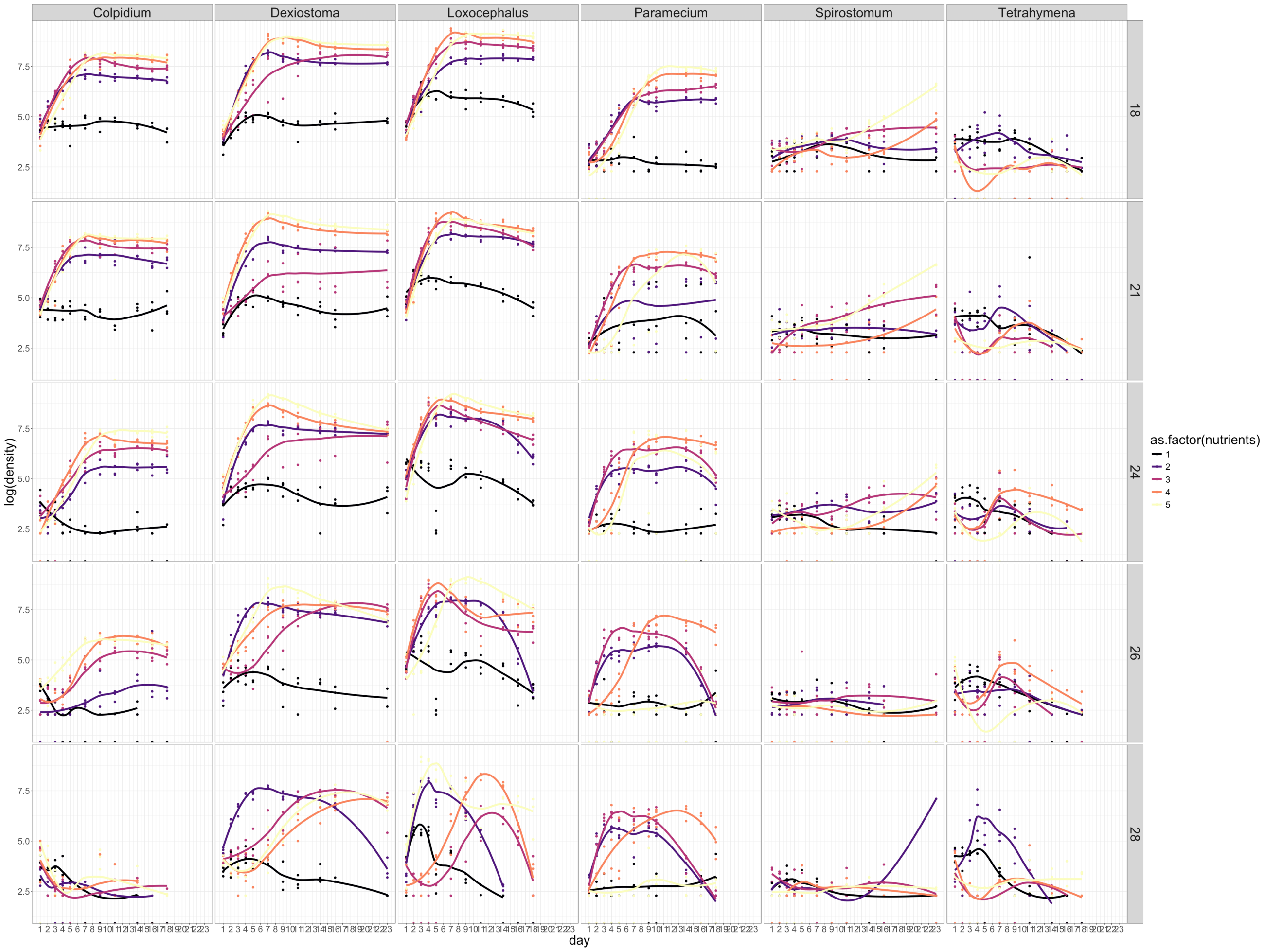


**Figure 1**:Time series of species densities across the treatments.

Calculate K and r. For some species in some environmental conditions, different lengths of the time series have been used to calculate intrinsic rate of growth

Visual inspection of exponential growth phase

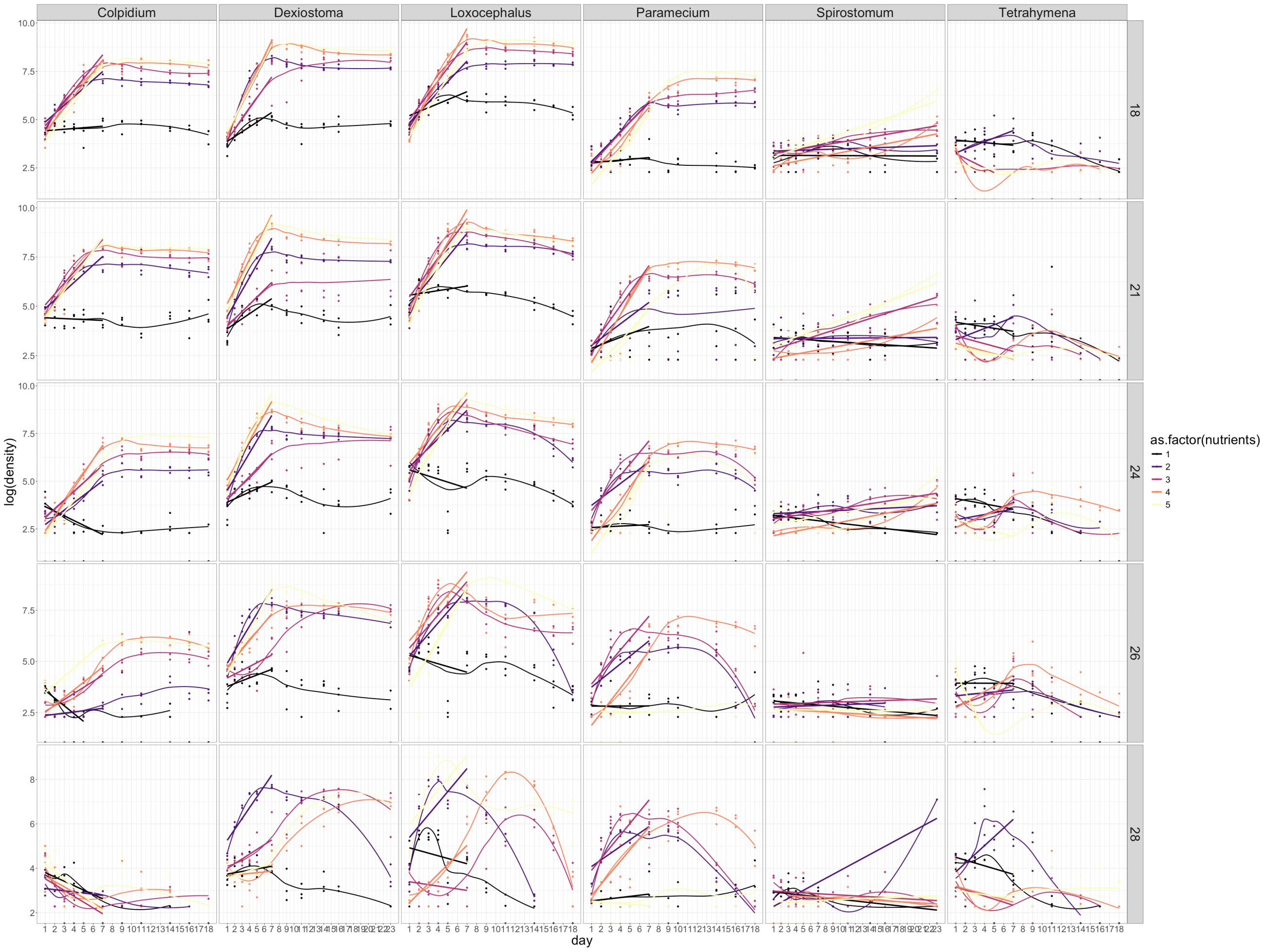


(#fig:reg_all)Species densities across the treatments with the regression lines used to calculate the intrinsic rate of growth (r).

### 1.1 GAMs to fit responde surface

We focus on the intrinsic rate of growth (r). This is because the intrisic rate of growth has been shown to be a better predictor of community temporal stability than carrying capacity (K) (Ross et al 2023) [https://besjournals.onlinelibrary.wiley.com/doi/full/10.1111/2041-210X.14087 (https://besjournals.onlinelibrary.wiley.com/doi/full/10.1111/2041-210X.14087)]. We are going to fit response surfaces using the calculated r for all species using GAMs. Then we are also going to use r as the species' trait to calculate potential response diversity, as well as response diversity when the trajecotry of the environmental change is known. The rationale is that r is likely of more relevance if the environmental change of interest occurs rapidly, since r provides information on a population's ability to rapidly bounce back after disturbance. In the upcoming experiment, we will have temperature fluctuating relatively fast, and so we decide now to focus on r.

Use GAMs to fit response surface of r and K

Create surface plots

We now check how the GAMs surfaces (r) look compared to the measured densities.

Checking predictions

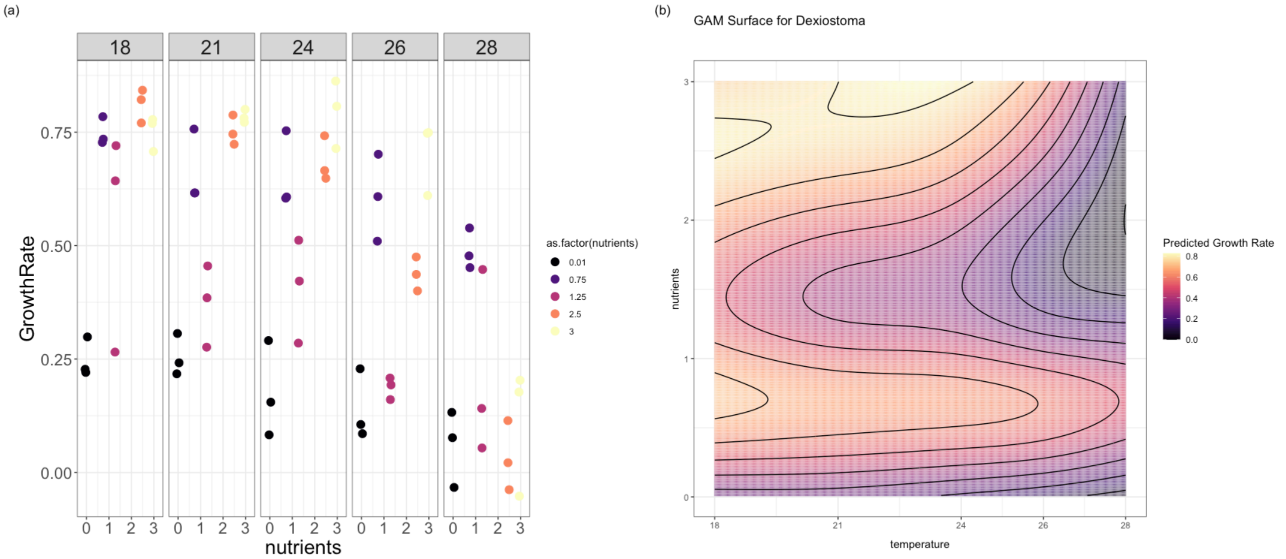


(#fig:surface_Dexi)Measured density values of Dexiostoma in the different treatments (a) vs fitted surface of growth rate (b).

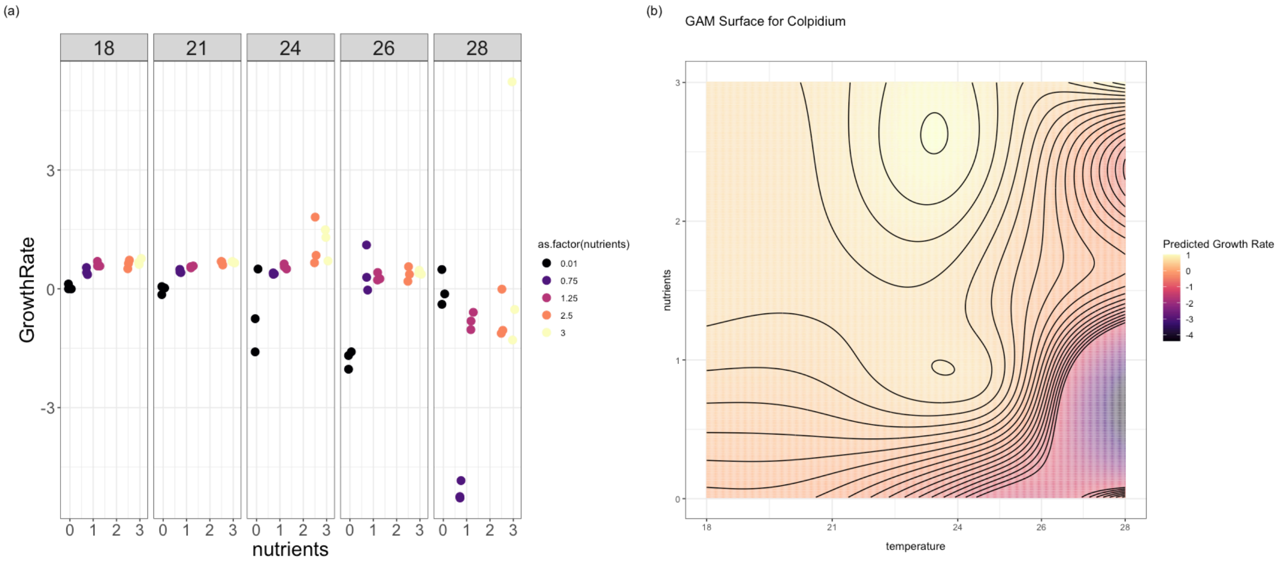


(#fig:surface_colp)Measured density values of Colpidium in the different treatments (a) vs fitted surface of growth rate (b).

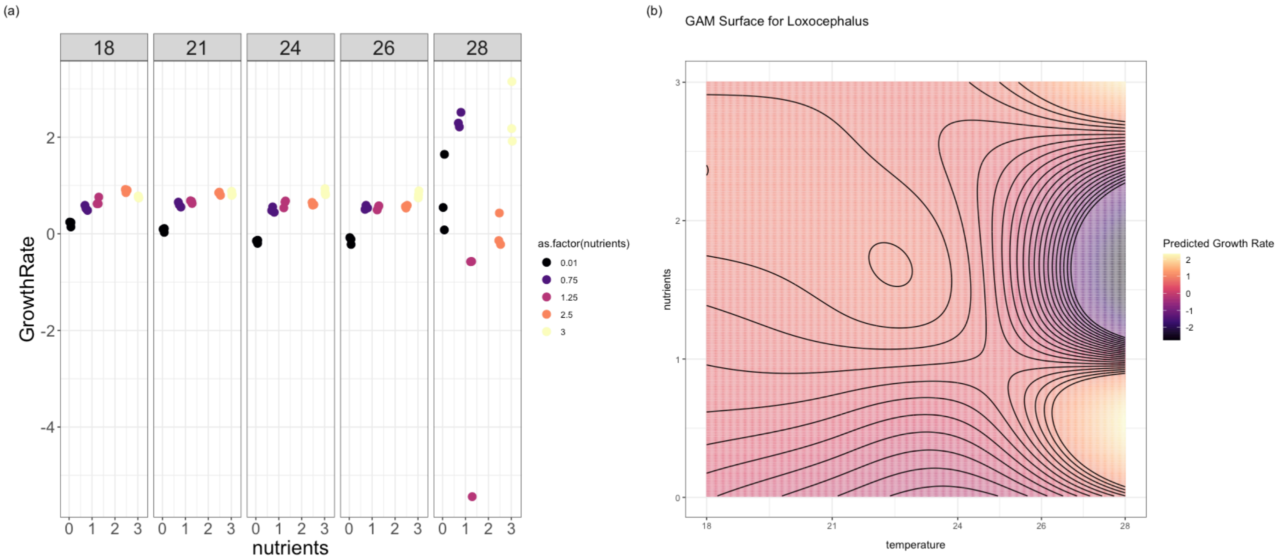


(#fig:surface_loxo)Measured density values of Loxocephalus in the different treatments (a) vs fitted surface of growth rate (b).

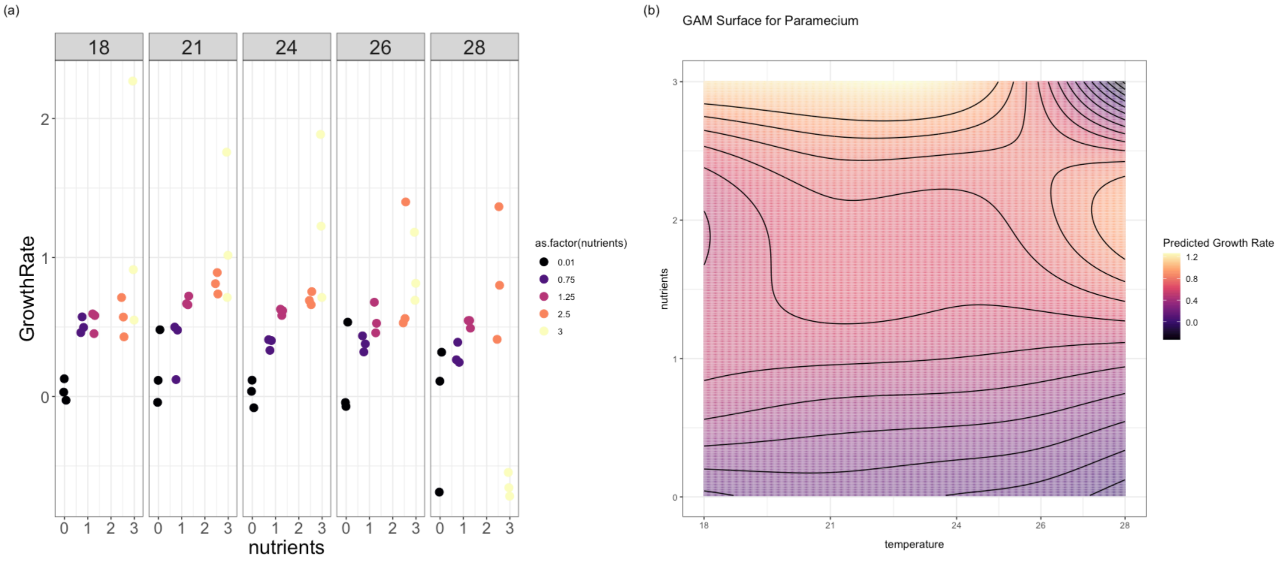


(#fig:surface_paramecium)Measured density values of Paramecium in the different treatments (a) vs fitted surface of growth rate (b).

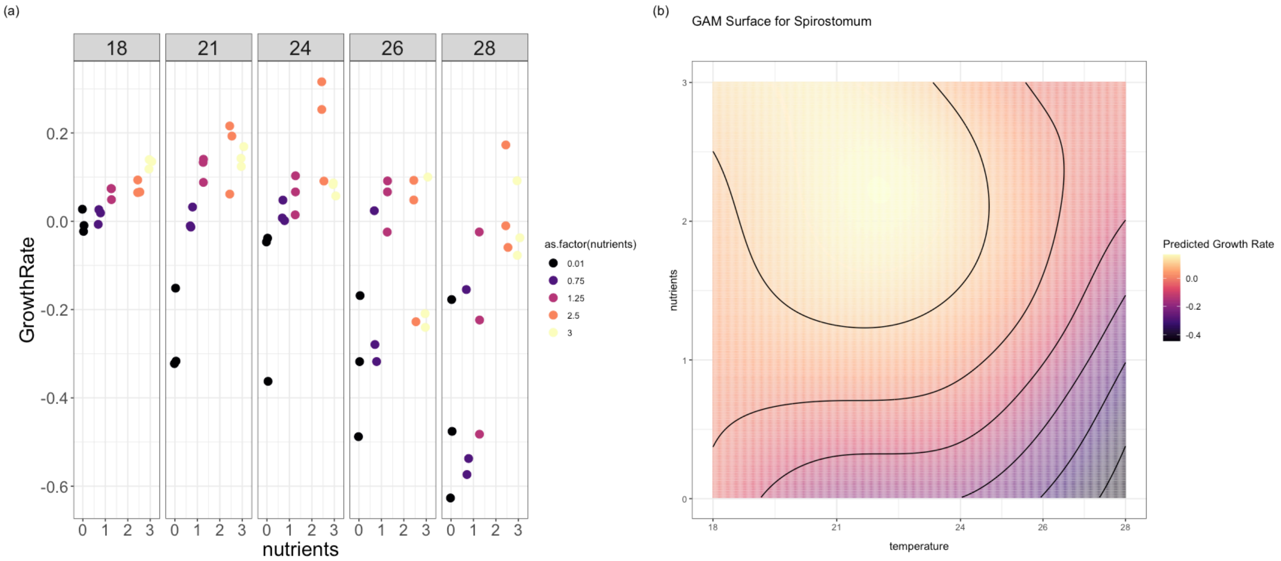


(#fig:surface_spiro)Measured density values of Spirostotum in the different treatments (a) vs fitted surface of growth rate (b).

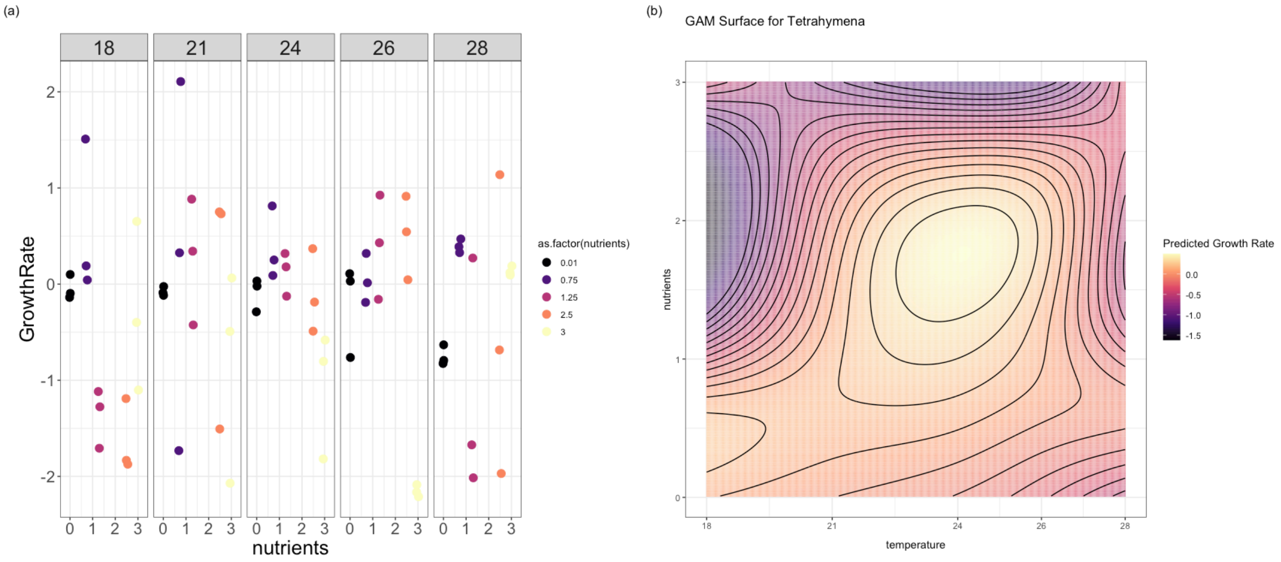


(#fig:surface_tetra)Measured density values of Tetrahymena in the different treatments (a) vs fitted surface of growth rate (b).

### 2 Response diversity with known direction of environmental change

Now we calculate RD with known trajecory of environmental change, i.e. the one we are going to apply in the experiment.

- Create a data set with environmental conditions that we may use in the experiment and calculate RD knowing the trajecotry of the environmental change.
- fluctuating temperature x 3 fixed nutrients = 9 treatments

#### 2.1 Temperature fluctuations

For the experiment we used three different temperature regimes, all with same magnitude but different mean temperatures: 18-21, 22-25, 25-28. Temperature stayed at lower end of the range for three days, and then transitioned over 24 h to the higher end, where it stayed for another three days before transitioning back. The experiment lasted for 60 days.

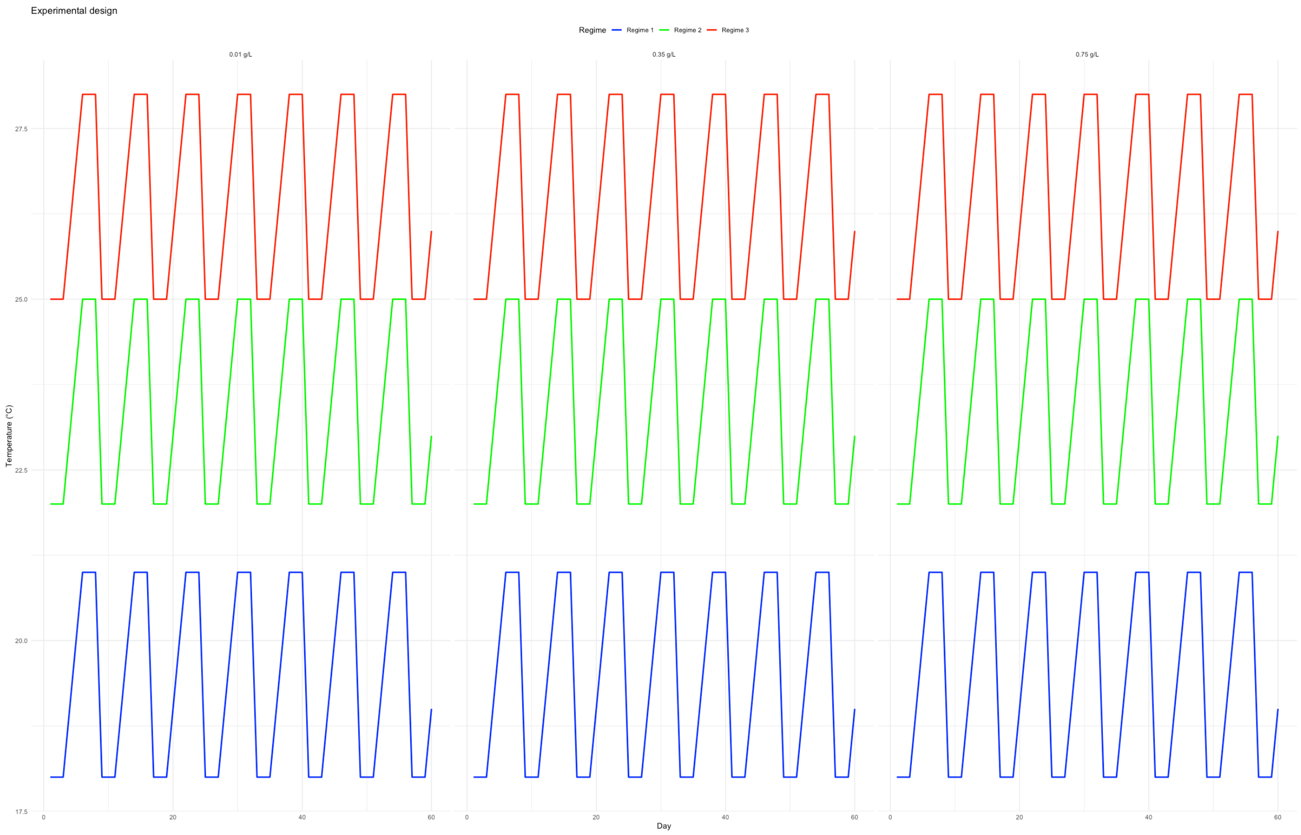


Figure 2.1: Visual representation of the experimental design.

#### 2.2 Get the slopes (change in species performance from Temperature 1 to Temperature 2 and vice versa)

### 2.3 Calculate diveregence for all possible compositions

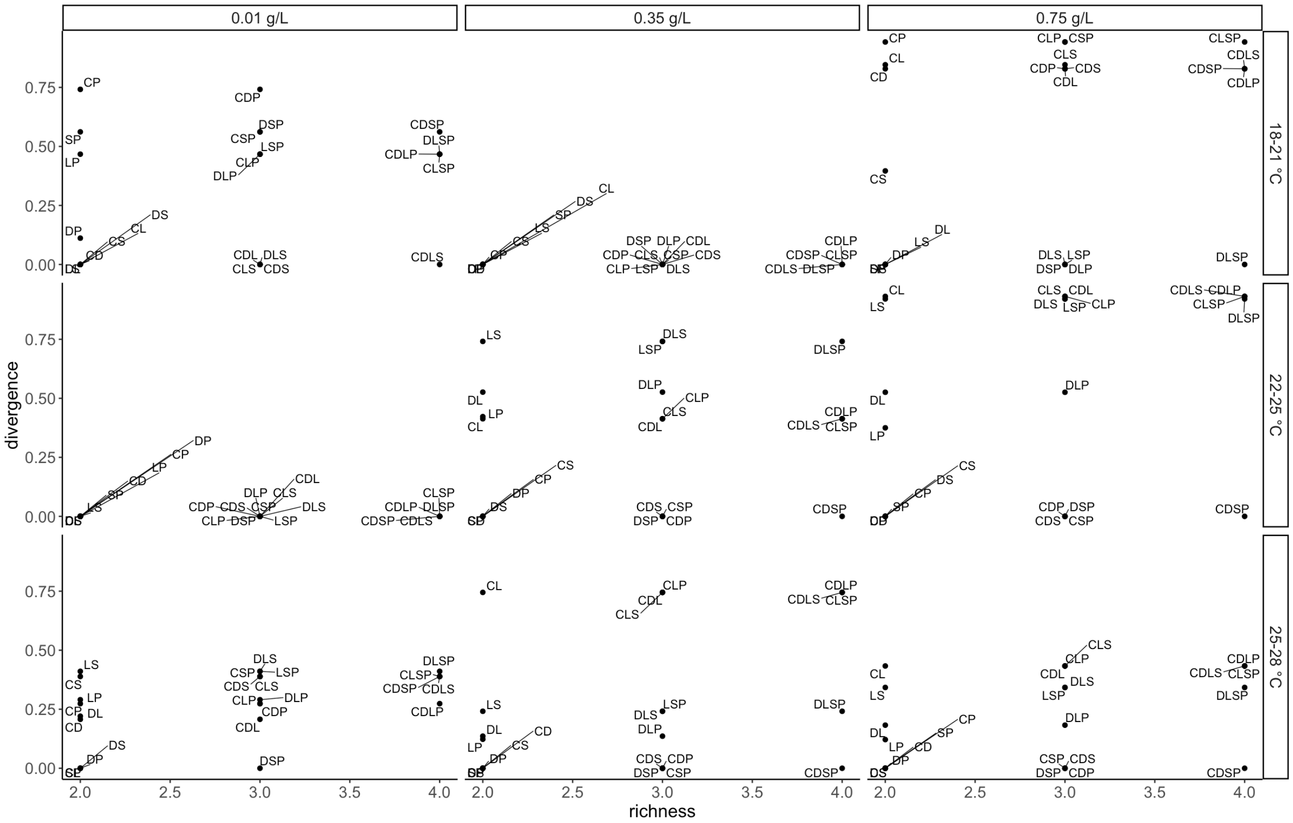


(#fig:communities_all)Community compositions that were used in the experiment to create, for each richness level, a gradient of response diversity. Letter represent different species: C = Colpidium, S = Spirostomum, D = Dexiostoma, P = Paramecium, L = Loxocephalus

### 2.4 Calculate diveregence

Communities employed in each environmental treatment were selected prior to the community experiment based on mono culture response experiments. While analyzing Community data using the BEMOVI R package, settings had to be changed (threshold and minimal particle size) to reduce processing time to a manageable level. To stay consistent between mono culture and community experiment (and to ensure correct species identification), we reanalyzed mono culture data. This resulted in slightly different response surfaces, causing some values of divergence to change. Therefore, the gradient of divergence has changed for some environmental treatments. In the plot below you can see the compositions used in the experiment and their response diversity for each enviromental treatment and richness level.

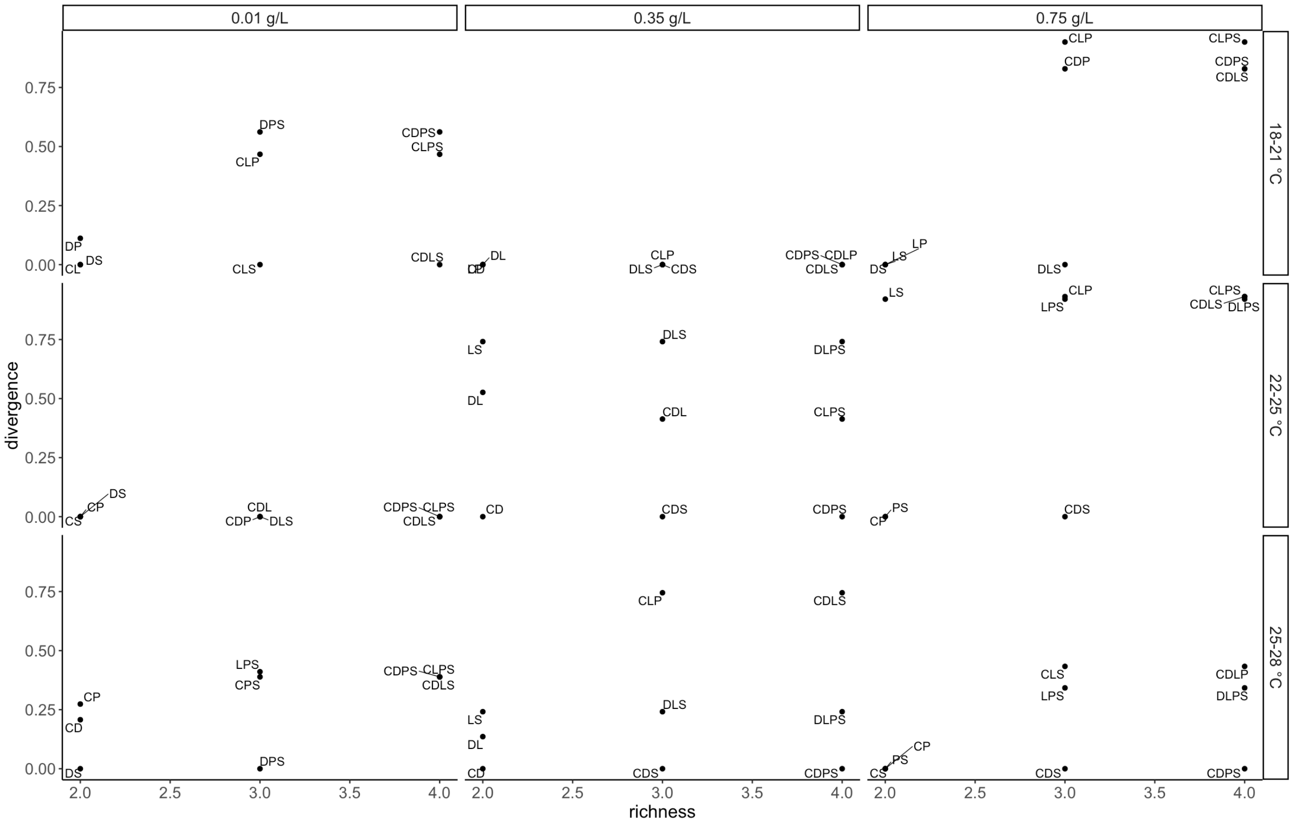


Figure 2.2: Community compositions that were used in the experiment to create, for each richness level, a gradient of response diversity. Letter represent different species: C = Colpidium, S = Spirostomum, D = Dexiostoma, P = Paramecium, L = Loxocephalus
