## Supplementary_info2 for "The imbalance of nature: The Role of Species Environmental Responses for Ecosystem Stability"

1 Introduction
2 Load data sets
3 Get time of extinction
4 Get average interaction coefficient
5 Get residual temperature and nutrients
6 Linear mixed effect model

### Species_interaction_analysis

Code

2025-01-27

### 1 Introduction

Competitive interactions have long been considered a key mechanism driving compensatory dynamics and promoting community temporal stability. Classic ecological theory posits that competition among species can reduce the dominance of any single species, thereby fostering asynchrony in population dynamics. This asynchrony, in turn, dampens fluctuations in total community biomass by ensuring that declines in one species are offset by increases in others. Such compensatory dynamics have been attributed to niche differentiation, resource partitioning, and trade-offs in species' environmental tolerances, which theoretically stabilize community functioning.

However, there is ongoing debate about the relative importance of competitive interactions versus fundamental species' responses to environmental variability in driving compensatory dynamics. While competitive interactions are thought to stabilize communities by promoting diversity and reducing synchrony, recent studies suggest that species' individual responses to environmental conditions may play a more dominant role. For example, variation in species' performance curves under changing environmental conditions can create compensatory patterns independent of direct interactions among species. This has led to a growing recognition that intrinsic environmental responses may outweigh competitive effects in determining community stability, particularly in systems experiencing strong environmental fluctuations.

In the present study, we investigated the relative contributions of competitive interactions and species' fundamental environmental responses to compensatory dynamics within experimental protist communities. By manipulating species richness and environmental conditions, we tested whether compensatory dynamics could be attributed primarily to competitive interactions or to the distribution of species' environmental responses. Our findings suggest that fundamental response diversity, rather than competitive interactions, was the primary driver of community temporal stability. Below, we provide a detailed analysis supporting this conclusion.

### 2 Load data sets

### 3 Get time of extinction

Coefficients were only considered for time points where species biomass did not drop below 1/10 of initial biomass

### 4 Get average interaction coefficient

Show

```
## `geom_smooth()` using formula = 'y ~ x'
```

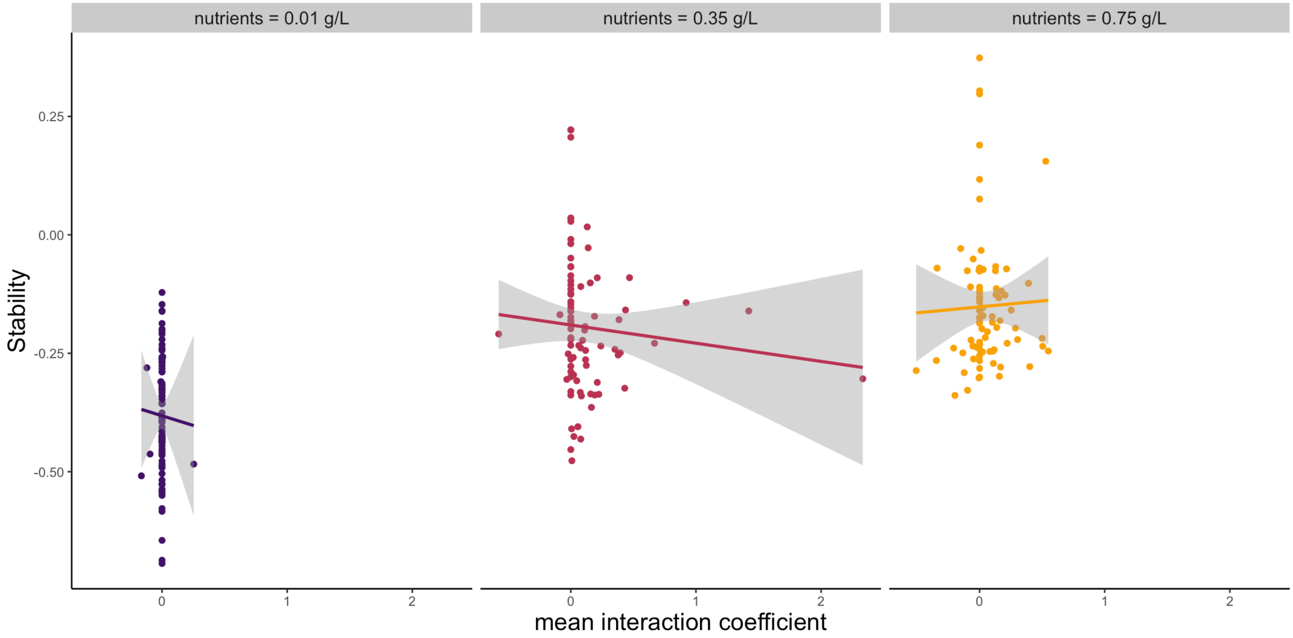


**Figure 1**: Relationship between mean interaction strength and stability divided by nutrient level.

Show

```
## `geom_smooth()` using formula = 'y ~ x'
```

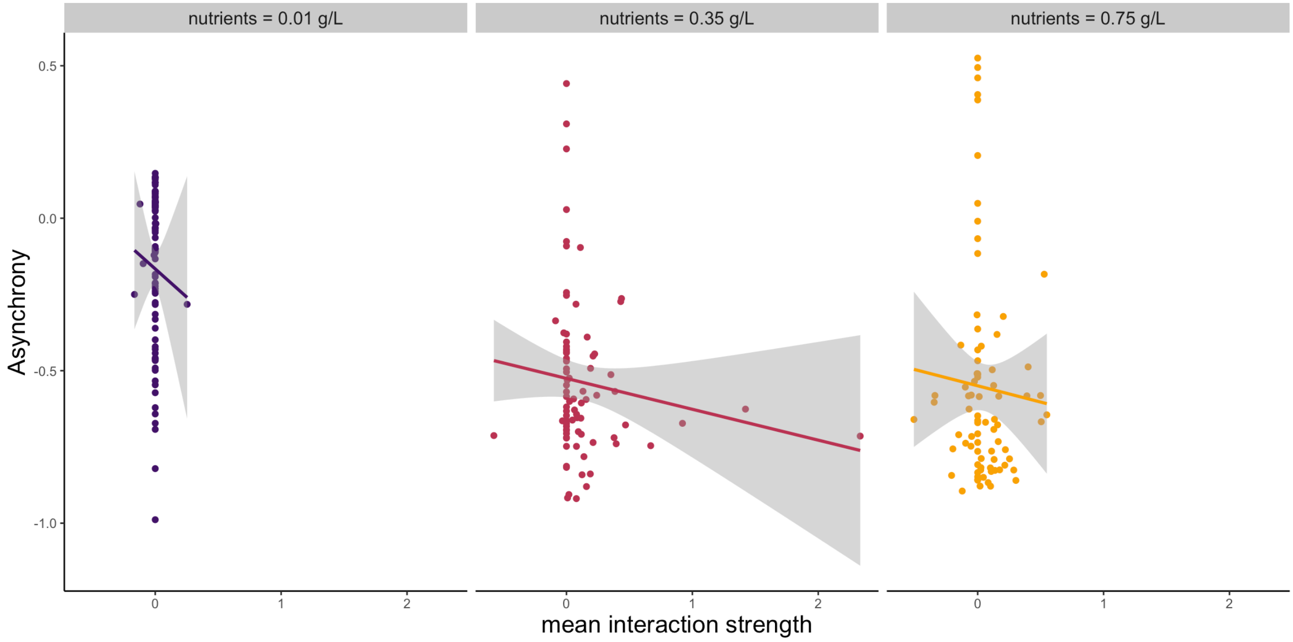


**Figure 2**: Relationship between mean interaction strength and asynchrony (Gross) divided by nutrient level.

Show

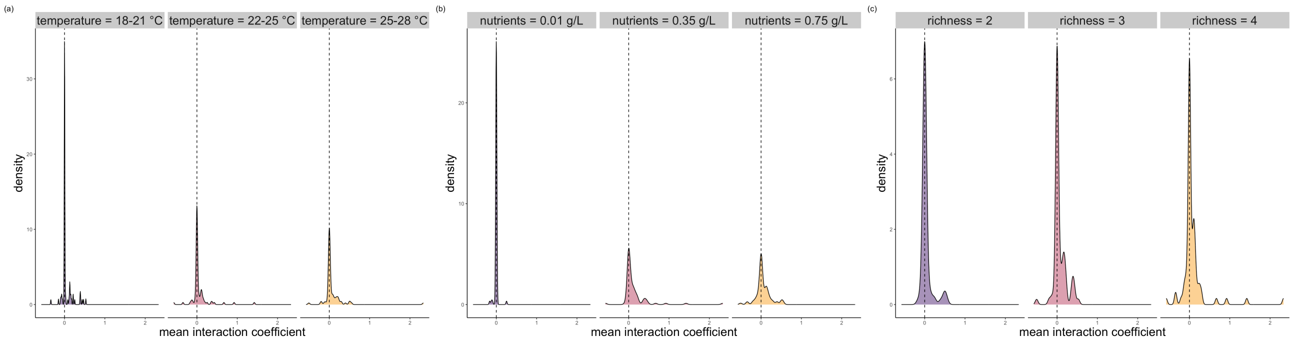


**Figure 2**: Distribution of mean interaction coefficient faceted by temperature, nutrients, and richness.

### 5 Get residual temperature and nutrients

### 6 Linear mixed effect model

The same model as in the main text, but with interaction coefficient as an additional explanatory variable. Negative interaction coefficients indicate competition while positive indicate facilitation.

**Table 2**: Linear mixed-effects model results for the effects of balance, species interactions, richness, and the residuals of temperature and nutrients on community stability. Estimates are presented with 95% confidence intervals and p-values.

Linear Regression Results

| Predictor | Estimate | 95% CI[1] | t-value | p-value |
|---|---|---|---|---|
| **Intercept** | -0.32 | -0.39, -0.24 | -8.88 | **<0.001** |
| **log10(balance_f)** | -0.08 | -0.09, -0.06 | -9.95 | **<0.001** |
| **richness** | | | | |
| richness3 - richness2 | -0.01 | -0.15, 0.12 | -0.27 | >0.9 |
| richness4 - richness2 | 0.01 | -0.15, 0.17 | 0.11 | >0.9 |
| richness4 - richness3 | 0.02 | -0.14, 0.18 | 0.32 | >0.9 |
| **resid_temp** | -0.10 | -0.12, -0.08 | -10.67 | **<0.001** |
| **resid_nut** | 0.25 | 0.21, 0.28 | 13.88 | **<0.001** |
| **mean_interaction** | 0.03 | -0.01, 0.07 | 1.34 | 0.2 |
| **resid_temp * resid_nut** | -0.09 | -0.15, -0.03 | -2.93 | **0.004** |

[1] CI = Confidence Interval

Mean interaction coefficient is not significant (est. = 0.03, p=0.2). No effect of interaction strength on stability was found.
